# Evolution of tunnels in α/β-hydrolase fold proteins – what can we learn from studying epoxide hydrolases?

**DOI:** 10.1101/2021.12.08.471815

**Authors:** Maria Bzówka, Karolina Mitusińska, Agata Raczyńska, Tomasz Skalski, Aleksandra Samol, Weronika Bagrowska, Tomasz Magdziarz, Artur Góra

## Abstract

The evolutionary variability of a protein’s residues is highly dependent on protein region and function. Solvent-exposed residues, excluding those at interaction interfaces, are more variable than buried residues whereas active site residues are considered to be conserved. The abovementioned rules apply also to α/β-hydrolase fold proteins - one of the oldest and the biggest superfamily of enzymes with buried active sites equipped with tunnels linking the reaction site with the exterior. We selected soluble epoxide hydrolases as representative of this family to conduct the first systematic study on the evolution of tunnels. We hypothesised that tunnels are lined by mostly conserved residues, and are equipped with a number of specific variable residues that are able to respond to evolutionary pressure. The hypothesis was confirmed, and we suggested a general and detailed way of the tunnels’ evolution analysis based on entropy values calculated for tunnels’ residues. We also found three different cases of entropy distribution among tunnel-lining residues. These observations can be applied for protein reengineering mimicking the natural evolution process. We propose a ‘perforation’ mechanism for new tunnels design via the merging of internal cavities or protein surface perforation. Based on the literature data, such a strategy of new tunnel design could significantly improve the enzyme’s performance and can be applied widely for enzymes with buried active sites.

**Author Summary:** So far very little is known about proteins tunnels evolution. The goal of this study is to evaluate the evolution of tunnels in the family of soluble epoxide hydrolases - representatives of numerous α/β-hydrolase fold enzymes. As a result two types of tunnels evolution analysis were proposed (a general and a detailed approach), as well as a ‘perforation’ mechanism which can mimic native evolution in proteins and can be used as an additional strategy for enzymes redesign.

## Introduction

Protein evolution mechanisms, and the factors determining protein evolution rate, have drawn attention in the past decades. Comprehensive studies regarding protein evolution resulted in a set of principles linking protein evolution with their structural and functional features. The most crucial assumption is that functionally important residues evolve at slower rates compared with the less important residues(1). Moreover, residues buried in the protein core and those on the protein surface were shown to have different substitution patterns(2), which may be related to different packing densities in the macromolecule(3). These findings provided the groundwork for various experimental techniques(4) and bioinformatic tools used intensively to carry out protein engineering(5, 6) to search for particular ancestral proteins(7–9), and to explore the evolution of enzyme functions within superfamilies(10). The distinction between residues that evolve more slowly or more quickly (i.e. conserved and variable residues, respectively) can be used to inform preselection of target regions for function or stability improvement, and in the design of smart libraries, while also providing explanations for unsuccessful attempts which resulted in dysfunctional or unstable mutants(11–15).

The amino acids comprising an enzyme’s catalytic site (regardless of its location) represent one of the most evident examples of conserved residues. In contrast, solvent-exposed residues which do not contribute to protein-protein or protein-ligand recognition are more variable, since they are not essential for either the enzyme’s function nor its structural stability(3,16,17). The evolutionary rate of secondary structure elements has also been investigated by several research groups. In a study by Sitbon and Pietrokovski(18), the authors suggest that, due to their regular repetitive structure, helices and strands might be more conserved than loops. On the other hand, Liu *et al.* showed that loops might tend to be evolutionary conserved since functional sites are overrepresented by loop-rich regions(19). However, other results suggest that β-sheet regions evolve more slowly in comparison to helical regions, and that random coil regions evolve the fastest(3,18,20,21).

Meanwhile, the results of site-directed mutagenesis experiments demonstrated that even mutations positioned relatively far from catalytic residues can attenuate an enzyme’s catalytic activity(22, 23). However, frequently distal mutations are fine-tuning the conformational ensembles of enzymes by evolutionary conformational selection(24, 25) but that approach can also modify the allosteric mechanism of an enzyme(26, 27), or its tunnel utilised to maintain ligands transport(28, 29). Growing evidence of a large number of tunnels in protein structures(28, 30) and their importance for an enzyme’s catalytic performance has led to the assumption that, while respecting evolutionary pressures, tunnels are generally preserved during protein evolution. So far, only a few individual studies have addressed this question. Evolutionarily preserved tunnels, or their parts, were reported in glutamine amidotransferases(31), carbamoyl phosphate synthetase(32), and histone deacetylases(33). In contrast, a faster rate of evolution was proposed for residues constituting gates in cytochromes(34).

Limited information about the variability of the tunnel-lining residues encouraged us to perform the first systematic study on the determination of a tunnel’s evolution in the soluble epoxide hydrolases (sEHs) family. We chose representative members of the sEHs due to three facts: i) that they belong to one of the oldest and the biggest enzymes superfamily - the α/β-hydrolases fold family(35–37), ii) that the crystal structures of different clade members (mammals, plants, fungi, and bacteria) were available, and iii) that sEHs catalyse the conversion of a broad spectrum of substrates and exhibit a diverse tunnel network in their structures. Such a tunnel network connects the conserved active site buried between the main and more structurally variable cap domain with the environment. We hypothesised that tunnels are conserved structural features equipped with variable parts, e.g. gates responsible for different substrate specificity in closely related family members. Additionally, we raised the following question: are there any mechanisms or schemes that can be adopted during protein engineering to mimic new tunnels’ appearance? Our results indicate that most tunnels in soluble epoxide hydrolases can be considered as conserved features, and we have proposed a “perforation” model that can be applied as a strategy for *de novo* tunnel design. Due to high structural similarity between members of α/β-hydrolases superfamily, our results could be expanded and applied into other superfamily members including acetylcholinesterase, dienelactone hydrolase, lipase, thioesterase, serine carboxypeptidase, proline iminopeptidase, proline oligopeptidase, haloalkane dehalogenase, haloperoxidase, epoxide hydrolase, hydroxynitrile lyase and others(38). We need to emphasise that since we analysed tunnels identified in relatively small protein structures with narrow tunnels (usually 1.0-2.0 Å), some processes leading to tunnel formation or modification cannot be covered. This includes long insertion or deletion, dimerization, or quaternary protein structure organisation.

## Results

For this study, we chose only the unique and complete structures of sEHs deposited in the Protein Data Bank (PDB)(39). Any structures with information missing about the positions of any of their amino acid residues could have provided bias, and therefore were excluded. The resulting selection of seven epoxide hydrolase structures represent the clades of animals (*Mus musculus*, msEH, PDB ID: 1CQZ, and *Homo sapiens*, hsEH, PDB ID: 1S8O), plants (*Solanum tuberosum*, StEH1, PDB ID: 2CJP), fungi (*Trichoderma reesei*, TrEH, PDB ID: 5URO), bacteria (*Bacillus megaterium*, bmEH, PDB ID: 4NZZ) and thermophilic enzymes collected in hot springs in Russia and China from an unknown source organism (Sibe-EH, and CH65-EH, PDB IDs: 5NG7 and 5NFQ, respectively).

### Model description and referential compartment evolutionary analysis

sEHs consist of two domains: the main domain, featuring eight β-strands surrounded by six α-helices; and the mostly helical cap domain, which sits atop the main domain. The cap domain is inserted between the strands of the main domain and is connected by an element called the NC-loop. The cap-loop is inserted between two helices of the cap domain(40). The active site of the sEHs is buried inside the main domain, and therefore the transportation of substrates and products is facilitated by tunnel (either singly or in a network)(29).

We performed an entropy analysis of the residues making up particular protein compartments with the use of the Schneider entropy metric implemented in the BALCONY package(41). As an input BALCONY requires multiple sequence alignment (MSA) and a list of residues building up particular compartments. We analysed the compartments listed in **Supplementary Table S1** (i.e. residues forming the active site; buried and surface residues; main and cap domains; NC-loop; cap-loop; and α-helices, loops, and β-strands). In order to determine the positions’ variability, we used Schneider entropy metric(42) calculated for each position in the MSA. To avoid bias and position-specific conservation scores we trimmed the MSA removing positions that did not correspond to the analysed proteins’ sequences. To evaluate the overall compartments’ variability we calculated the difference between the median distances of positions of the proteins‘ compartments and the remaining positions of the trimmed MSA (**Figure 1, Supplementary Table S2,** see also **Methods** section for the description of the MSA trimming). Negative values of the difference between median distances of the selected proteins’ compartments and the trimmed MSA (**Supplementary Table S2**) indicate compartments with lower variability, and positive values indicate compartments with higher variability in comparison to the remaining positions in the trimmed MSA. For quantitative statistical analysis, we compared the calculated Schneider entropy values of these compartments with the remaining positions of the trimmed MSA using the Epps-Singleton test(43).

**Figure 1.**
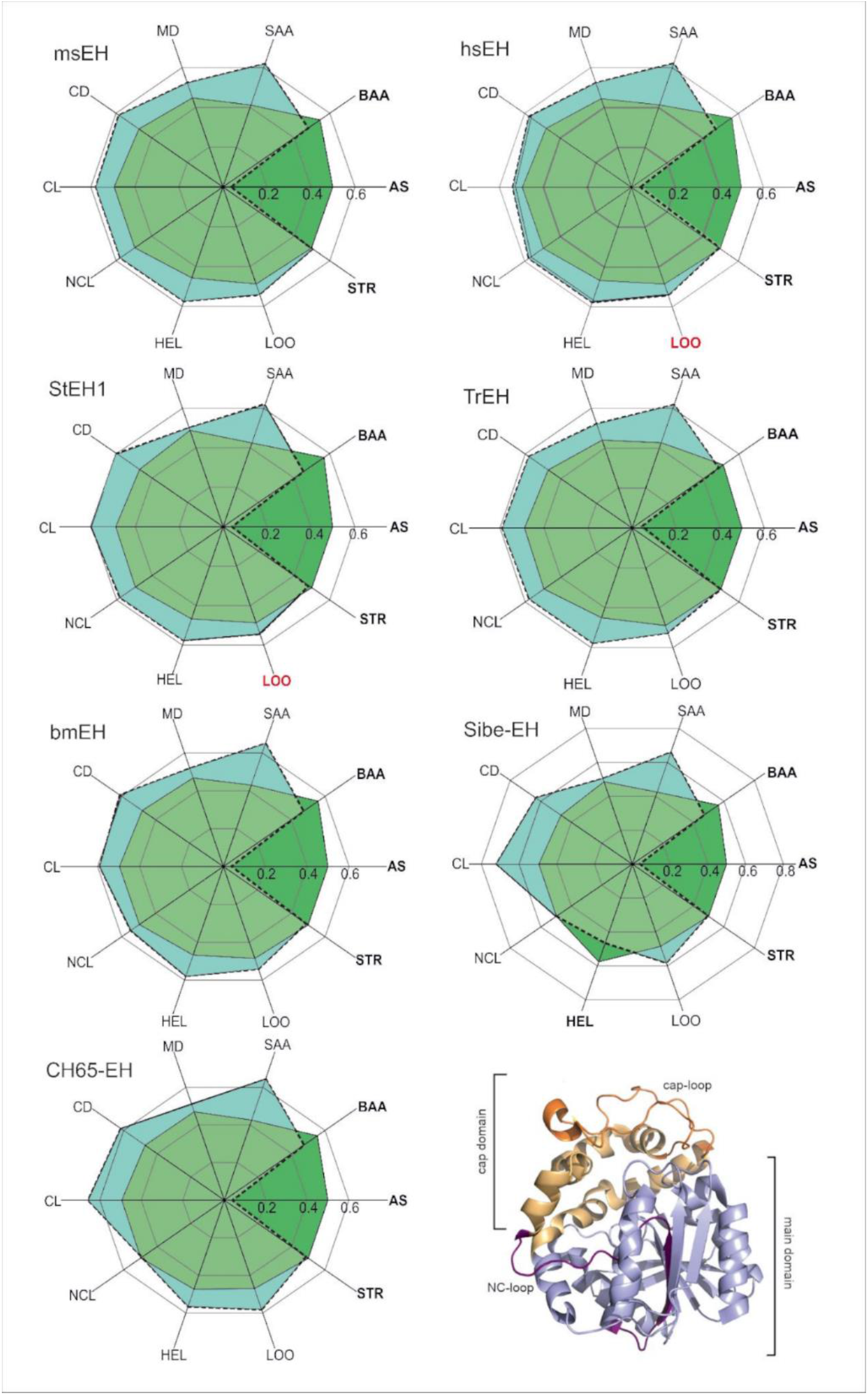
Radial plot of the median entropy values of referential compartments (green) and the remaining positions of the trimmed MSA (turquoise). When the median entropy values of the components cover the median entropy values of the trimmed MSA, it means that the particular compartment is more conserved than the remaining positions of the MSA (dark green). The compartments considered as conserved are written in bold. The MSA contained 1455 sequences and 419 positions. Figure represents data shown in **Supplementary Table S2**. All pairwise differences (except for loops (LOO) in hsEH and StEH1, marked red) are statistically significant (Epps-Singleton test). In the bottom right corner, a schematic representation of the analysed structure-specific comportments is provided. Abbreviations: AS – active site; BAA – buried amino acids; SAA – surface amino acids; MD – main domain; CD – cap domain; CL – cap-loop; NCL – NC-loop; HEL – helices; LOO – loops; STR – strands.

Based on the obtained differences in median distances and the results of the Epps-Singleton test, the active site residues were classified as conserved, i.e. with lower entropy scores in comparison to the remaining positions in the MSA. The surface residues (classified as solvent-exposed residues according to the NetSurfP server(44)) were classed as the most variable. Entropy analysis showed that the variability of the buried residues was significantly lower than the variability of the surface residues (**Figure 1, Supplementary Table S2**). These results are in agreement with the general findings mentioned previously(3,16,45). With regards to the structural elements specific to EHs, all compartments (main domain, cap domain, cap-loop, and NC-loop (except for the NC-loop in CH65-EH)) were classified as variable among all the selected sEHs. In all analysed proteins, α-helices and loops were also classified as variables (however, in the case of hsEH and StEH1 the information about the variability of loops was not statistically significant). In all analysed proteins, except for msEH, β-strands were found to be conserved which stays in agreement with the work of Sitbon and Pietrokovski(18) (**Figure 1, Supplementary Table S2**).

### Tunnel identification and comparison

We identified tunnels providing access to the active site using a geometry-based approach implemented in CAVER software(46) for both crystal structures and in molecular dynamics (MD) simulations, and then compared their geometries (for details see the **Methods** section). CAVER software identified between three and nine tunnels in the analysed crystal structures. Those tunnels were then compared with the tunnels identified during MD simulations to find their corresponding counterparts (**Supplementary Table S3**), based on the similarity of their tunnel-lining residues (for more details see the **Methods** section). We marked all identified tunnels according to their localisation within the epoxide hydrolase’s domains as was shown in our previous work(47). We identified tunnels passing through three regions of the sEH structure: i) the main domain (marked as Tg, Tm, Tback, and Tside), ii) the cap domain (marked as Tcap), as well as iii) the border between the cap and main domains (marked as Tc/m).

We identified seven tunnels in the main domain, six in the cap domain, and three at the border between those domains (**Figure 2**). It should be pointed out that the Tc/m tunnel was identified as multiple tunnels by CAVER (Tc/m1, Tc/m2, and Tc/m3). This issue is related to the asymmetric shape of the Tc/m tunnel, which makes it difficult to classify in a geometry-based approach (**Supplementary Figure S1**).

**Figure 2.**
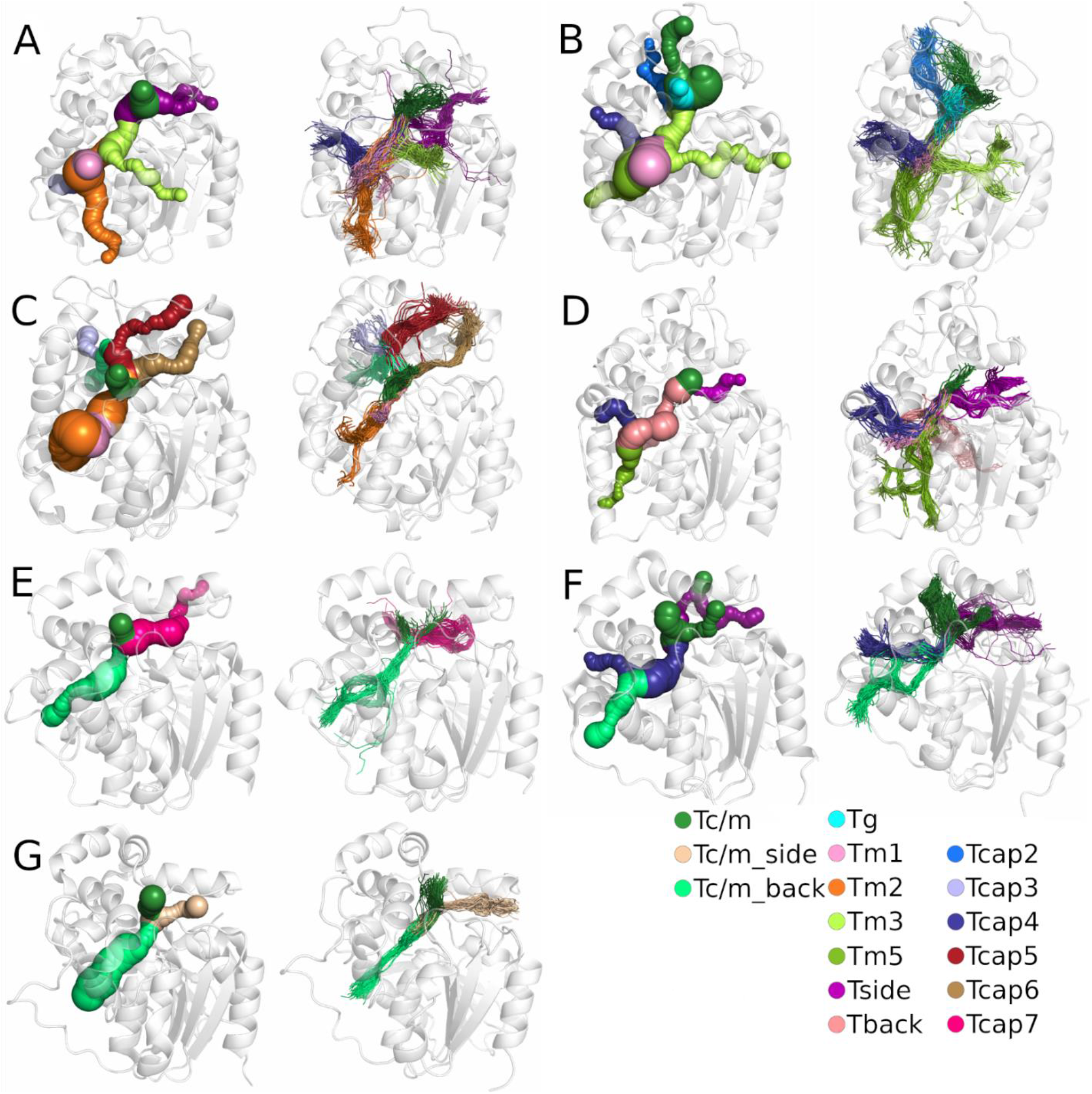
Comparison of tunnels identified in the crystal structure (left) and the results after molecular dynamics (MD) simulation (right) for each system: A) *M. musculus* soluble epoxide hydrolase (msEH), B) *H. sapiens* soluble epoxide hydrolase (hsEH), C) *S. tuberosum* soluble epoxide hydrolase (StEH1), D) *T. reesei* soluble epoxide hydrolase (TrEH), E) *B. megaterium* soluble epoxide hydrolase (bmEH), and thermophilic soluble epoxide hydrolases from an unknown source organism F) Sibe-EH, and G) CH65-EH. Protein structures are shown as white transparent cartoons. Matching tunnels are marked with the same colour as spheres (in crystal structures) and lines (in MD simulations).

Closer analysis of the tunnels identified in crystal structures and during MD simulations by CAVER showed that the tunnels identified in crystal structures are well-defined; however, their parts located closer to the protein surface are, in some cases, coiled. For most tunnels identified during MD simulations, the interior parts of tunnels were well-defined, whereas the tunnels’ mouths were widely distributed on the protein surface. Such an observation might suggest that those regions are tightly packed and/or lined by bulky residues which can change their conformation to open/close a particular tunnel.

### Tunnel evolutionary analysis

In the case of sEHs, tunnels can perform several distinct functions: i) transport and positioning of substrates and products, ii) control of the solvent access to the catalytic cavity, and iii) transport of catalytic water. Only those tunnels which maintain at least one of those functions can undergo evolutionary pressure. As we confirmed during the referential compartments’ evolutionary analysis, surface residues are more variable than buried residues. Indeed, **Figure 3** shows protein structures coloured according to Schneider entropy values, where thin blue lines represent regions with lower entropy, and yellow thick lines represent regions with higher entropy values. We also coloured the identified tunnels according to their frequency of detection (i.e. based on the number of frames in which they were identified) in MD simulations (darker = more frequent). The overall position of the tunnels was similar among all the protein structures; however, there were large differences concerning their frequency during the MD simulations. Cross-sections of these structures suggest that the protein core is composed of residues with lower variability (lower entropy values), whereas the tunnel mouths, located at the protein surfaces, are surrounded by residues of both higher and lower variability (higher and lower entropy values, respectively).

**Figure 3.**
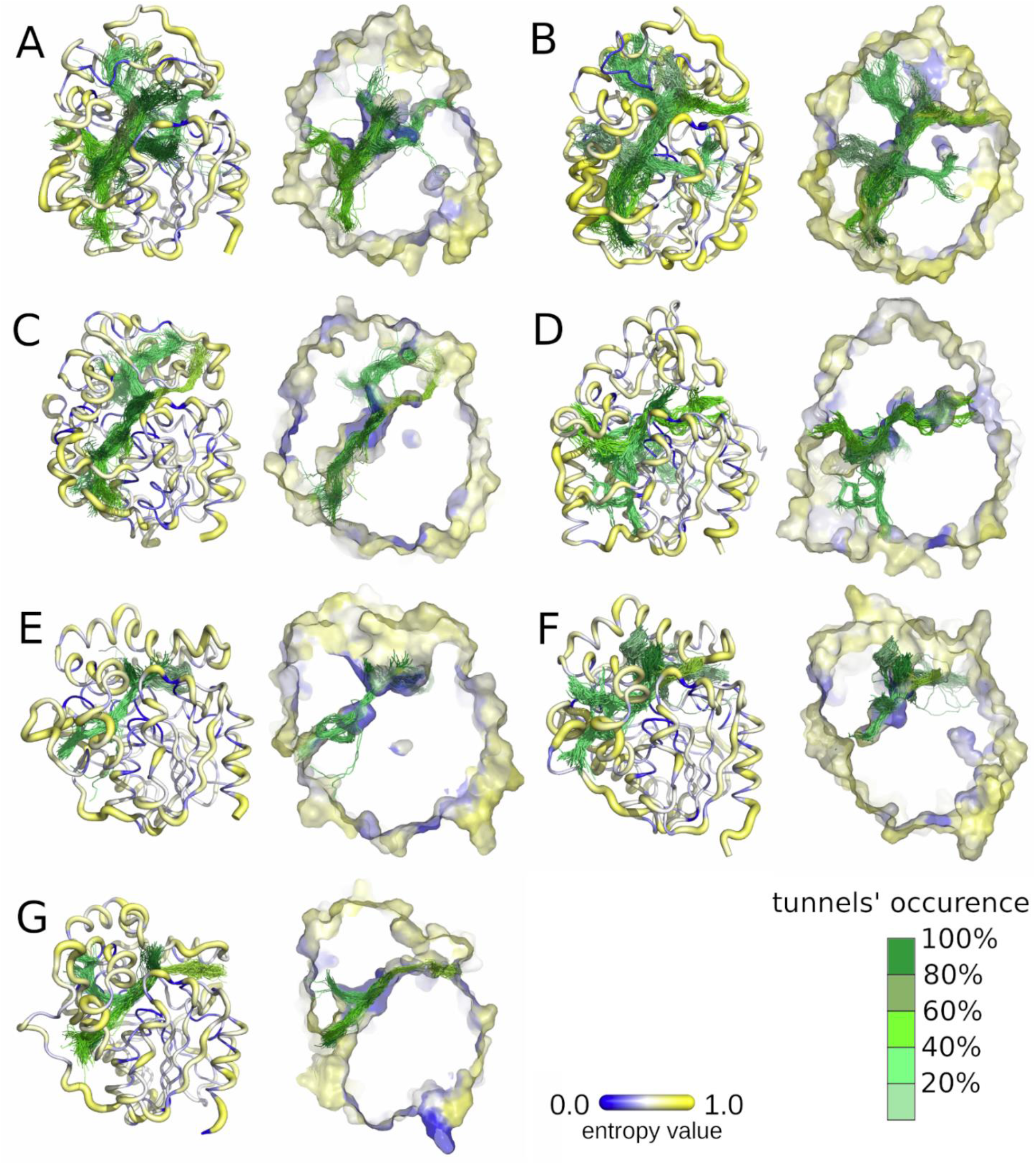
Visualisation of the entropy score of each protein residue (right), and frequency of tunnels identified with CAVER during molecular dynamics (MD) simulations (left) for each system: A) *M. musculus* soluble epoxide hydrolase (msEH), B) *H. sapiens* soluble epoxide hydrolase (hsEH), C) *S. tuberosum* soluble epoxide hydrolase (StEH1), D) *T. reesei* soluble epoxide hydrolase (TrEH), E) *B. megaterium* soluble epoxide hydrolase (bmEH), and thermophilic soluble epoxide hydrolases from an unknown source organism F) Sibe-EH, and G) CH65-EH. Protein residues are shown according to their entropy score: low values of entropy are marked as thin blue lines and higher values as thick yellow lines. Tunnel centerlines are coloured according to the frequency of their occurrence during MD simulations (the tunnels occurrence was calculated based on the numbers of the MD simulation frames in which the tunnel was identified; 100% means that the tunnel remained open in all 50,000 MD simulation frames): dark green indicates the most frequently identified tunnels, and light green those very rarely identified. The right side of each pair shows cross-sections of protein surfaces coloured according to the entropy score of each amino acid residue.

We identified the residues lining these particular tunnels during the MD simulations. During MD simulations, the protein is not a rigid body and the residues gain some level of flexibility, which may cause the opening and closing of identified tunnels. Moreover, due to the residues’ movements, the identified tunnels may branch (either near the active site, in the middle of the tunnel, or near the surface). Since we observed many cases of tunnels branching near the surface, the list of identified tunnel-lining residues may be overrepresented by the surface residues. Therefore, we decided to perform an entropy analysis of: i) all tunnel-lining residues; ii) surface tunnels-lining residues; and iii) tunnel-lining residues without the surface residues. An evolutionary analysis of the tunnel-lining residues without the surface residues is presented in **Figure 4**. Analysis was performed using the same procedure as in the case of the referential compartments. Complete lists of tunnel-lining residues are shown in **Supplementary Tables S5–S11**. A detailed analysis of the sEHs tunnels is shown in **Supplementary Figures S3–S9**. Median distances of all analysed proteins are listed in **Supplementary Table S12**.

**Figure 4.**
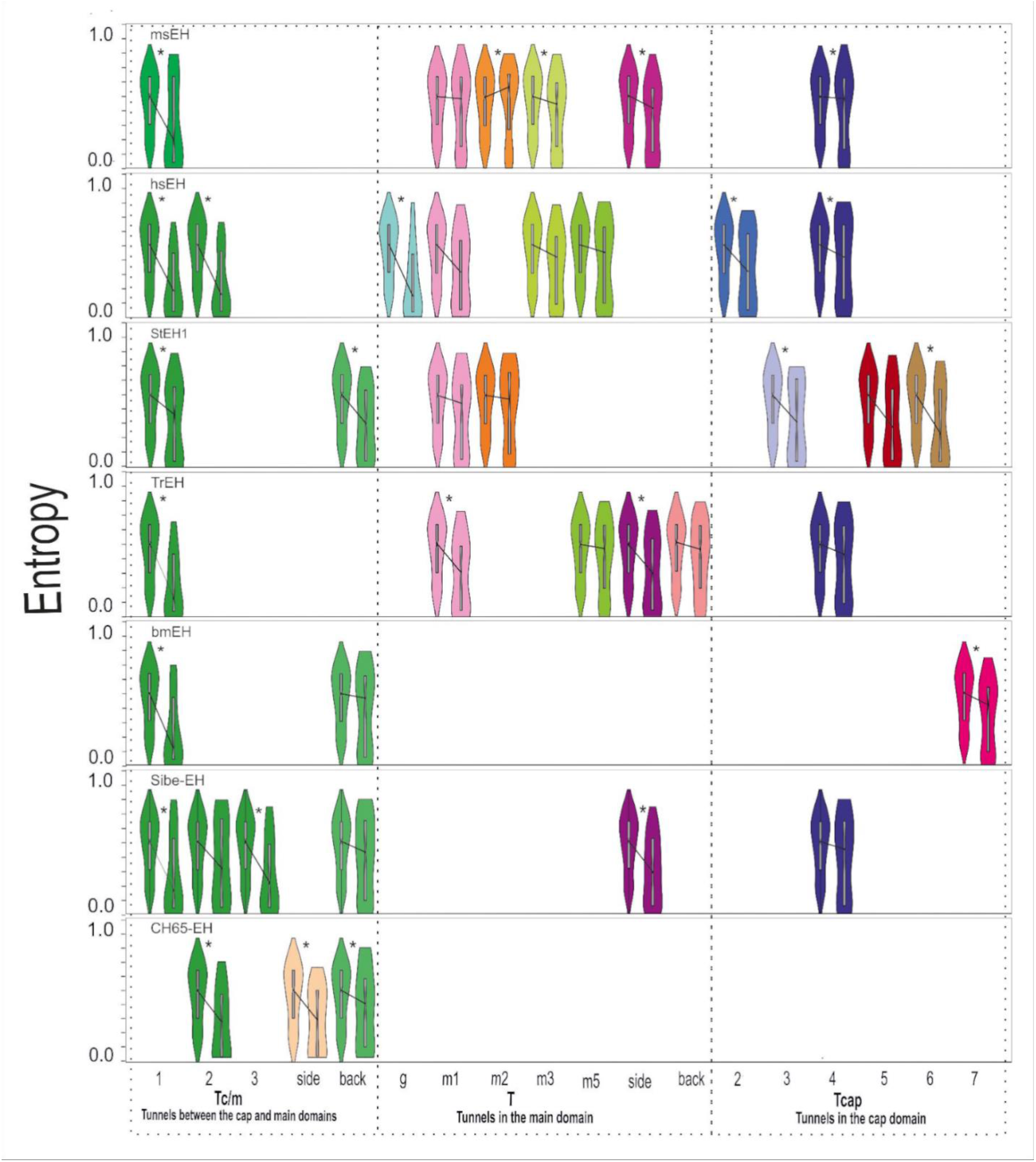
Distribution of the entropy values and median entropy values of tunnel-lining residues without the surface residues, and the remaining positions of the trimmed MSA (violin plots), for all analysed soluble epoxide hydrolase structures. Figure represents the data shown in **Supplementary Table S12**. Statistically significant pairwise differences in median distances are marked by a star (*).

Based on the median distances between all tunnel-lining residues and the remaining residues’ positions in the MSA, we concluded that almost all analysed tunnels should be considered as conserved. Following exclusion of the surface residues from the tunnel-lining residues, differences in median values decreased indicating that the conserved character of tunnels comes from the buried residues (**Figure 4, Supplementary Table S12**). It is clear that the surface tunnel-lining residues generally reach higher entropy values than the other analysed tunnel-lining residues (**Supplementary Figures S3–S9**).

Presented violin plots (**Figure 4**) provide insight into tunnel’s residues entropy distribution. To perform that, the right violin shape from each pair has to be analysed. For example, in bmEH, the distribution of the entropy values among residues creating Tc/m1 tunnel shows a triangle-like shape with a wide base of residues with low entropy values, which corresponds to the prevailing contribution of conserved residues. In contrast, the distribution of the entropy values among residues lining the Tc/m_back tunnel resembles a rectangle or even hourglass-like shape which means that both variable and conserved residues build that tunnel. Thus, analysis of the shape of the violin plots provides descriptive information about the variability of the residues creating each tunnel. The differences between violin plots for all tunnel-lining residues, and those with excluded surface residues, clearly confirm the variable character of tunnels’ entries.

### Detailed analysis of selected tunnels

The violin plots provide information about the general variability of the tunnel-lining residues, but do not give insight into the location of the variable/conserved residues along the tunnel. To analyse that we have performed a more advanced analysis. We selected three different tunnels which were identified in three different sEHs. The Tc/m tunnel of hsEH and the Tm1 tunnel of StEH1 represent the most commonly identified tunnels, and the Tc/m_back tunnel of bmEH represents an interesting case of a tunnel which already was engineered. The entropy values of the tunnel-lining residues are presented in **Supplementary Table S13**.

As we pointed out elsewhere(47) the Tc/m tunnel whose mouth is located between the main and cap domains can be seen as an ancestral tunnel created during cap domain insertion and preserved in nearly all epoxide hydrolases. In hsEH this tunnel (**Figure 5, panel A**) has an average length of ∼13.3 Å. It was open during 59% of the simulation time, with an average bottleneck radius of 1.6 Å, reaching a maximum of 2.7 Å. It is lined by residues with both low and high values of entropy, which makes the overall entropy distribution nearly flat (when surface residues are included) or exponential (when surface residues are excluded) which corresponds to the hourglass-like and triangle-like shape of the violin plot, respectively. The majority of variable residues is located close to the surface or at the interface between the cap and main domains. Close inspection of the tunnel revealed also a highly variable residue (i.e. with higher entropy value) – F497 (Schneider entropy value 0.7946) – located approximately in the middle of the tunnel and situated between two less-variable residues (i.e. with lower entropy values) – D496 (Schneider entropy value 0.0336), from the active site, and V498 (Schneider entropy value 0.4713). The F497 residue might act as a molecular gate(48) since its position in several other crystal structures differs substantially, and was identified as a surface residue (**Supplementary Figure S10**).

**Figure 5.**
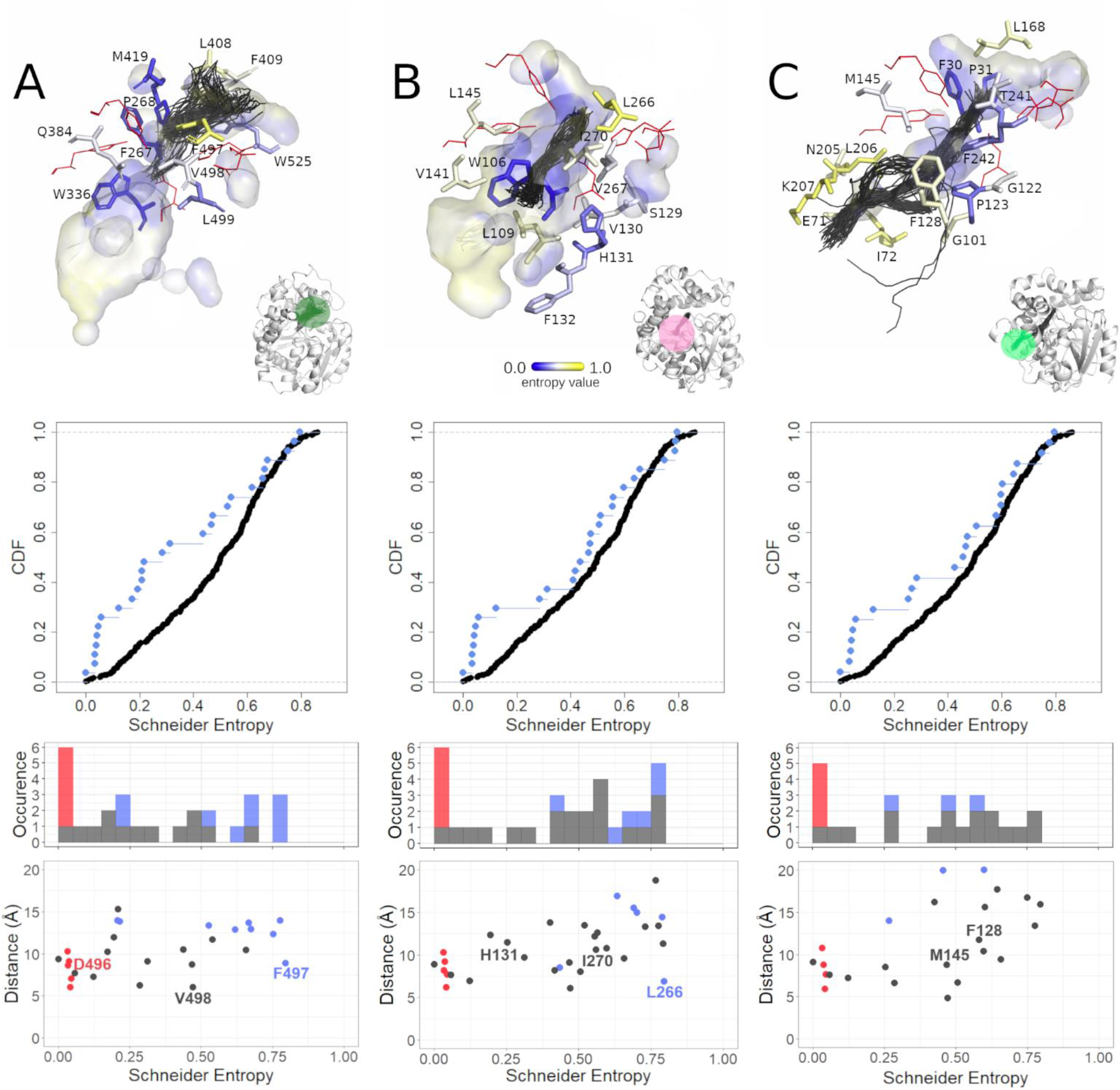
Analysis of the selected tunnels of soluble epoxide hydrolases (sEHs). A) The Tc/m tunnel of the *H. sapiens* soluble epoxide hydrolase (hsEH) structure, B) the Tm1 tunnel of the *S. tuberosum* soluble epoxide hydrolase (StEH1) structure, and C) the Tc/m_back tunnel of the *B. megaterium* soluble epoxide hydrolase (bmEH). Each panel consists of three parts: top section - close-up of tunnel residues. Residues are coloured according to entropy score. For the sake of clarity, less-frequently detected amino acid residues were omitted, and those creating the active site are shown as red lines. The active site cavity is shown as the interior surface, and the representative tunnel detected during molecular dynamics (MD) simulations as centerlines; middle section - cumulative distribution function (CDF) of entropy score for the tunnel-lining residues without the surface residues (cyan dots) and corresponding counterpart (black dots); and bottom section – scatterplot of the tunnel residues’ entropy values relative to distance from the geometric centre of the α carbons of the enzyme, along with a marginal histogram of entropy value counts in respective intervals. Scatterplot points as well as histogram counts grouped into classes based on residue classification (active site – red; surface residues – blue; buried – grey).

The Tm1 tunnel of StEH1 is the shortest identified in this structure (**Figure 5, panel B**). Similar tunnels were identified in three other analysed sEHs: msEH, hsEH, and TrEH. The tunnel mouth is located in the main domain, near the NC-loop and hinge region. A close inspection of this tunnel revealed that it was ∼13 Å long on average, and was always open during MD simulation. It had an average bottleneck radius of 1.9 Å, with a potential to increase up to 3.1 Å. The analysis of the violin plots suggests overrepresentation of the variable residues (reversed triangle-like shape of the violin plot, when surface residues are included), and nearly flat distribution of entropy values (hourglass-like violin shape, when surface residues are included). The majority of the tunnel-lining residues showed relatively high entropy values, while the residues with lower entropy values were located in proximity to the active site. In our previous analysis of StEH1(49), we identified three residues, namely P188 (Schneider entropy value 0.5117) (not shown in Figure 5), L266 (Schneider entropy value 0.7946), and I270 (Schneider entropy value 0.5594), as potentially useful during protein engineering. Here we present that those residues are also variable, which may suggest that their substitution might not affect protein stability. Interestingly, approximately in the middle of the tunnel length, a less-variable H131 (Schneider entropy value 0.2524) residue was also identified.

The last example is the Tc/m_back tunnel of bmEH (**Figure 5, panel C**) which was already engineered by Kong *et al.*(50). This tunnel was identified as a third tunnel during MD simulation and had an average length of 26.7 Å. It was open only for 18% of the simulation time, with an average bottleneck radius of 1.0 Å, and the potential to increase up to 1.8 Å. The mouth of this tunnel was located in the main domain. Both violin plots (with surface residues included and excluded) show a similar hourglass-like shape. Close inspection of the tunnel revealed that residues with lower entropy values contributed to the binding cavity inside the main domain, while residues with higher entropy were located in the area surrounding a deep pocket on the protein surface. We also found two residues, namely F128 (Schneider entropy value 0.5798) and M145 (Schneider entropy value 0.4678), which had lower entropy values than their neighbours. Those residues were successfully modified to create a novel tunnel leading to the bmEH active site, allowing conversion of bulky substrates(50).

## Discussion

In our study, we focused on sEHs, which are enzymes belonging to the α/β-hydrolase superfamily. Members of this superfamily share a barrel-like scaffold of eight anti-parallel β-strands surrounded by α-helices with a mostly helical cap domain sitting on top of the entrance to the active site(45), which seems to be also the oldest(51) and most stable(52) fold used by one the largest groups of enzymes(53). Structural and evolutionary analyses of EHs have been reported systematically(40,47,54–56), providing valuable insights into their structural and functional features. In our work, we first assessed the system-specific compartments described previously by Barth *et al.*, such as the main and cap domains, the NC-loop, and the cap-loop, along with secondary structure elements such as strands, helices, and loops. Based on an alignment of 95 EH sequences, three available crystal structures, and several homology models, they showed that the main and cap domains are conserved, while the NC-loop and cap-loop are variable(40).

Here, we analysed an alignment of 1455 EH sequences and additionally performed an in-depth analysis of the seven complete crystal structures representing different clades (animals, plants, fungi, and bacteria). By calculating the difference between median distances of Schneider entropy values of a selected compartment and the remaining positions of the trimmed MSA – we were able to determine the variability of each compartment. The calculated median distances for all analysed compartments confirmed well-known observations: active sites comprised highly conserved residues, with greater variability exhibited by surface residues than by buried residues(3,16,45). Our results were also consistent with the work of Barth *et al.*(40), showing that the cap-loop and NC-loop should be considered as variable features. However, in contrast to their work, for such a large set of sequences, the whole main and cap domains were considered variable. In all analysed proteins, α-helices, and loops were found to be variable (**Supplementary Table S2**), while β-strands were found to be conserved in all analysed proteins, except for msEH. Such a tendency was shown previously for other systems elsewhere(18). Further, since we were able to identify structural compartments of the seven sEHs analysed, observations regarding the modularity of EHs are still applicable(37).

The main aim of our analysis was to perform what was, to our knowledge, the first systematic analysis of the evolution of tunnels in a large family of sEHs. Therefore, our results can be applied mostly to the EHs, and – with some minor adjustments – to other members of the α/β-hydrolases superfamily. We identified multiple tunnels of different sizes and shapes, located in three different regions: the cap and main domains, as well as at the border between those domains. We hypothesised that tunnels are conserved structural features equipped with variable parts, such as gates responsible for different substrate specificity profiles in closely related family members. This hypothesis was based on two assumptions: i) that the surface residues are more variable in comparison to the buried residues, and ii) that access to the active site cavity should be preserved to sustain the catalytic activity of the enzyme. Our results confirmed both assumptions. Moreover, we identified the Tc/m tunnel which was present in all analysed sEHs, and is located in the border between the cap and main domains. The cap domain is thought to be a result of a large insertion into the α/β-hydrolase main domain(45, 47). Both domains interact, creating a hydrogen bond network(57); they co-evolved to preserve access to the buried active site while also ensuring the flexibility required for transport of the substrate and the products(58). Most of the residues with lower entropy values in the cap domain face the main domain. This finding confirms previously presented information about the main and cap domains’ relative flexibility(40,59–62).

We also proposed two ways of the analysis of the tunnel residues variability. The violin plots allow analysis of the contribution of variable and conserved residues, which provides a general overview of each tunnel. They also allow assessment of the variability of a particular compartment relative to the remaining positions of the MSA (as shown in **Figure 1**). The scatterplots (similar to those in **Figure 5, panel A-C**) provide detailed insight and can be used to draw further conclusions regarding the distribution of entropy values of tunnel-lining residues along an analysed tunnel. They can also be used to identify the most variable and conserved tunnel-lining residues. In general, after excluding the active site and surface residues, the analysed examples (**Figure 5**) show three cases of entropy distribution among tunnel-lining residues: i) the flat distribution of the entropy values (**Figure 5, panel A**); ii) the overrepresentation of residues with higher entropy values (**Figure 5, panel B**); and iii) the quasi-sigmoidal distribution (**Figure 5, panel C**; most of the residues have values of the entropy in the range of 0.25–0.7).

Our results confirmed the conserved character of the tunnels. Moreover, we found that even conserved tunnels can be lined with more variable residues, located not only at the surface (tunnels’ entry). Close inspection of the Tc/m tunnel of hsEH allowed us to detect variable S412 and F497 residues (Schneider scores 0.618 and 0.795, respectively), among which phenylalanine was observed to be the most flexible amino acid, and which was even observed in a different conformation in crystal structures (**Supplementary Figure S10**). This indicates a potential role for F497 as a gate, controlling access through this tunnel(48). On the other hand, the Tc/m tunnel is also lined by more conserved residues, such as the highly conserved substrate-stabilising tyrosine located in the cap domain (Y466 in hsEH, Schneider score 0.0323)(63, 64).

Analysis of the variability of particular amino acid positions could be used in the search for feasible key amino acids (hot-spots)(65). More variable positions might be considered as favourable locations for the introduction of mutations. Such residues can be detected even for the shortest tunnels, and have already been shown to enable fine-tuning of enzyme properties(66). For example, the Tm1 tunnel of StEH1 is lined with several variable residues which may have a role to play in the fine-tuning of the enantioselectivity of that enzyme(67). Such a strategy is acknowledged as one of the most likely to succeed, since it does not significantly disturb protein activity and stability, and the different locations of hot-spots along the transport pathway may enable modification of geometric/electrochemical constraints, thus contributing to the enzyme selectivity.

In our other study, we showed a relationship between a tunnel’s shape and location, and the enzyme’s function(47). Thus, the evolution of the tunnel network can be considered as an additional mechanism that allows the enzyme to adapt and catalyse the conversion of different substrates. Mimicking such a process could provide a straightforward strategy for enzyme re-engineering. As we pointed out above, the insertion of the cap domain has created the buried active site cavity and the Tc/m tunnel ensuring access to that cavity. This tunnel can be considered as an ancestral tunnel and it seems to be well-preserved among nearly all sEHs family members. However, the insertion of large fragments into existing structures appears to be a high-risk strategy. Based on our results, we can suggest a much easier approach that can be used for tunnel network redesign.

### Perforation mechanism of the tunnel formation

The observed entropy values of tunnel-lining residues usually range from 0.25 to 0.7 (**Supplementary Table S13**). As we showed, the scatterplots can be used to identify the most variable and conserved residues. Variable residues are considered potentially safe hot-spots for single-point mutations(65). We can imagine that new tunnels providing access to the protein interior can appear as a result of a “perforation” *via* a mutation occurring: i) in the surface layer of protein or ii) at the border of large cavities affecting surface integrity (**Figure 6**). Such a process can be easily mimicked and adopted for enzyme modification. We showed(47) that, in some cases, tunnels behave more like a series of small cavities which are rarely open. In the case of such tunnels, a mutation resulting in a permanently open cavity might be a driving force for future tunnel widening and modulation of selectivity or activity of enzymes, or otherwise provide additional regulation of activity.

**Figure 6.**
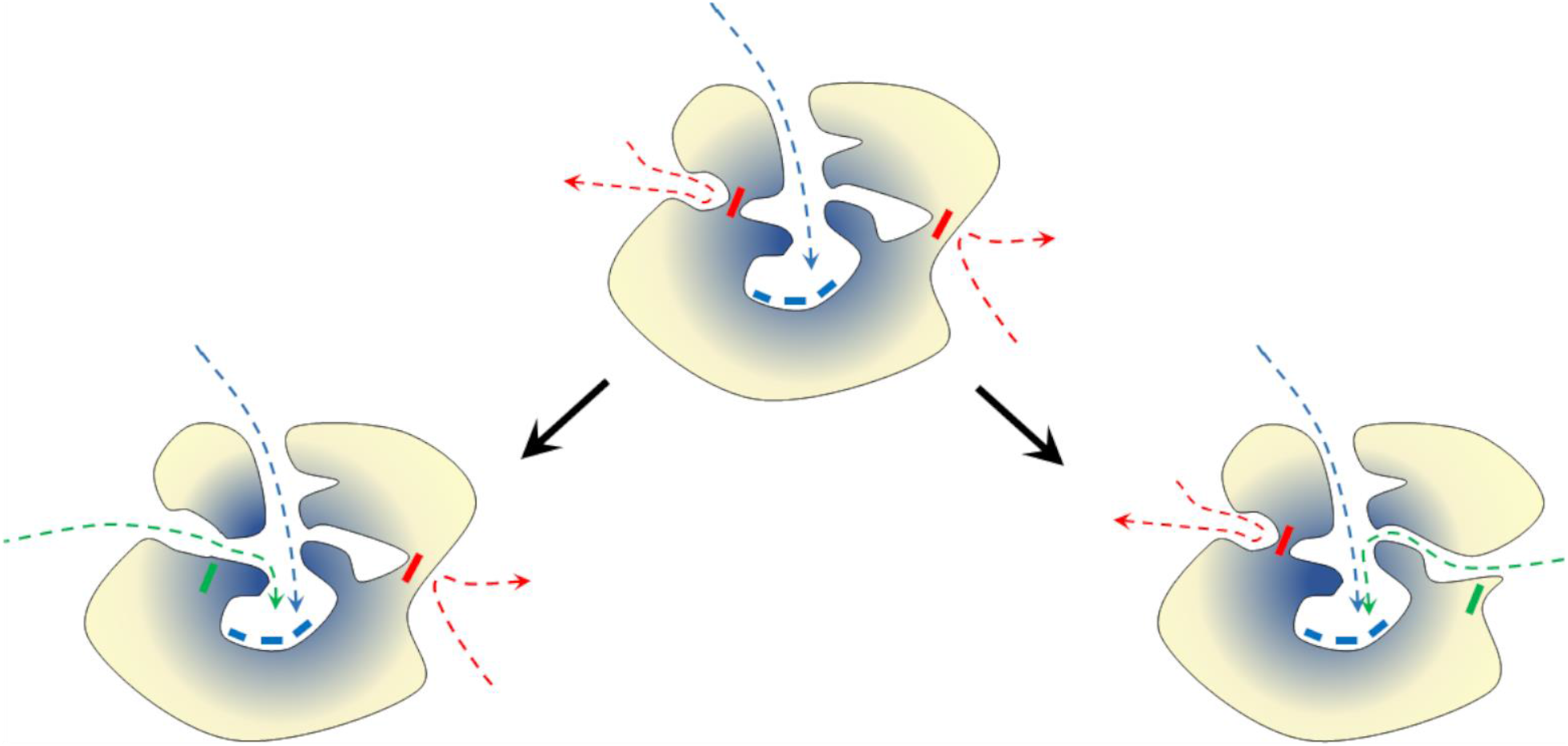
Schematic representation of the ‘perforation’ model of protein tunnel evolution. The ancestor protein (middle) and two modification pathways leading to a new enzyme by merging internal cavities (left) or by surface perforation (right). Yellow – variable residues; blue – conserved residues. Boxes represent residues: blue – conserved active site residues, red – potentially mutable (variable) residue, and green – mutated residue. Arrows represent pathways leading to the active site: blue – actual pathway, red – potential novel pathway, and green – novel open pathway.

The appearance of a new tunnel, resulting from a single-point mutation, *via* the proposed perforation mechanism provided significant freedom and flexibility for α/β-hydrolases to modify their activity and selectivity. Since the mechanism for hydrolysis performed by the sEHs involves deprotonation of the nucleophile in the hydrolysis step (proton shuttling) and water attack, it requires precise transport of water molecules. New tunnels could significantly improve the enzyme’s performance by separating the substrate/products transport pathways from water delivery tracks.

A perfect example of the mimicking of the proposed surface perforation model is the transformation of the Tc/m_back tunnels of the bmEH shown by Kong *et al.*(50) in which they turned a substrate inaccessible tunnel into an accessible one in order to improve the enzyme’s functionality. As we showed here these two residues whose substitution to alanine led to the opening of a side tunnel, improving the activity of bmEH upon α-naphthyl glycidyl ether, had higher entropy values than their neighbours. This work also led us to a hypothesis that mutations of such variable residues could also appear spontaneously and may drive the evolution of the active site accessibility *via* surface perforation and/or joining of internal cavities. Identification of such residues which are prone to cause such an effect might easily be adopted as part of protein reengineering processes. These conclusions are supported by the observations of Aharoni *et al.*(68), who noticed that most mutations affecting protein functionality (mostly activity and selectivity) were located either on the protein surface or within the active site cavity. Indeed, the investigation of long and narrow tunnels, not obviously relevant at first glance during protein engineering, should be regarded as a strategy for new pathway creation, as illustrated by Brezovsky *et al.* in their *de novo* tunnel design study which resulted in the most active dehalogenases known so far(69). Dehalogenases are closely related to sEHs and belong also to the α/β-hydrolases superfamily, thus further supporting the rationality of our approach.

The tunnels described in our findings which we consider conserved provided substantial information about their origin, and about the evolution of enzymes’ families more broadly. On the other hand, our results suggest that after the ancestral occlusion of the active site, the further evolution of α/β-hydrolases may be driven by perforation of either the surface or of the internal cavities, which mostly comprised variable residues. Tunnels themselves can be equipped with both conserved residues, which are potentially indispensable for their performance, as well as highly variable ones, which can be easily used for fine-tuning an enzyme’s properties. Such hotspots can be easily identified using the approach presented here.

## Methods

### Workflow

Evolutionary analysis was divided into two parts: system-specific compartment analysis, and tunnel analysis. Prior to those analyses, the positions of the residues that contribute to compartments and tunnels needed to be mapped in an MSA comprising sequences of epoxide hydrolases. The identified residues were then used as input for an evolutionary analysis using the BALCONY software(41). Tunnels were identified by CAVER software(46) in both crystal structures and during MD simulations and then compared with each other to find their corresponding counterparts. Finally, the tunnel-lining residues, the surface tunnel-lining residues, and the tunnel-lining residues without surface residues were used for the evolutionary analysis using BALCONY software (**Figure 7**).

**Figure 7.**
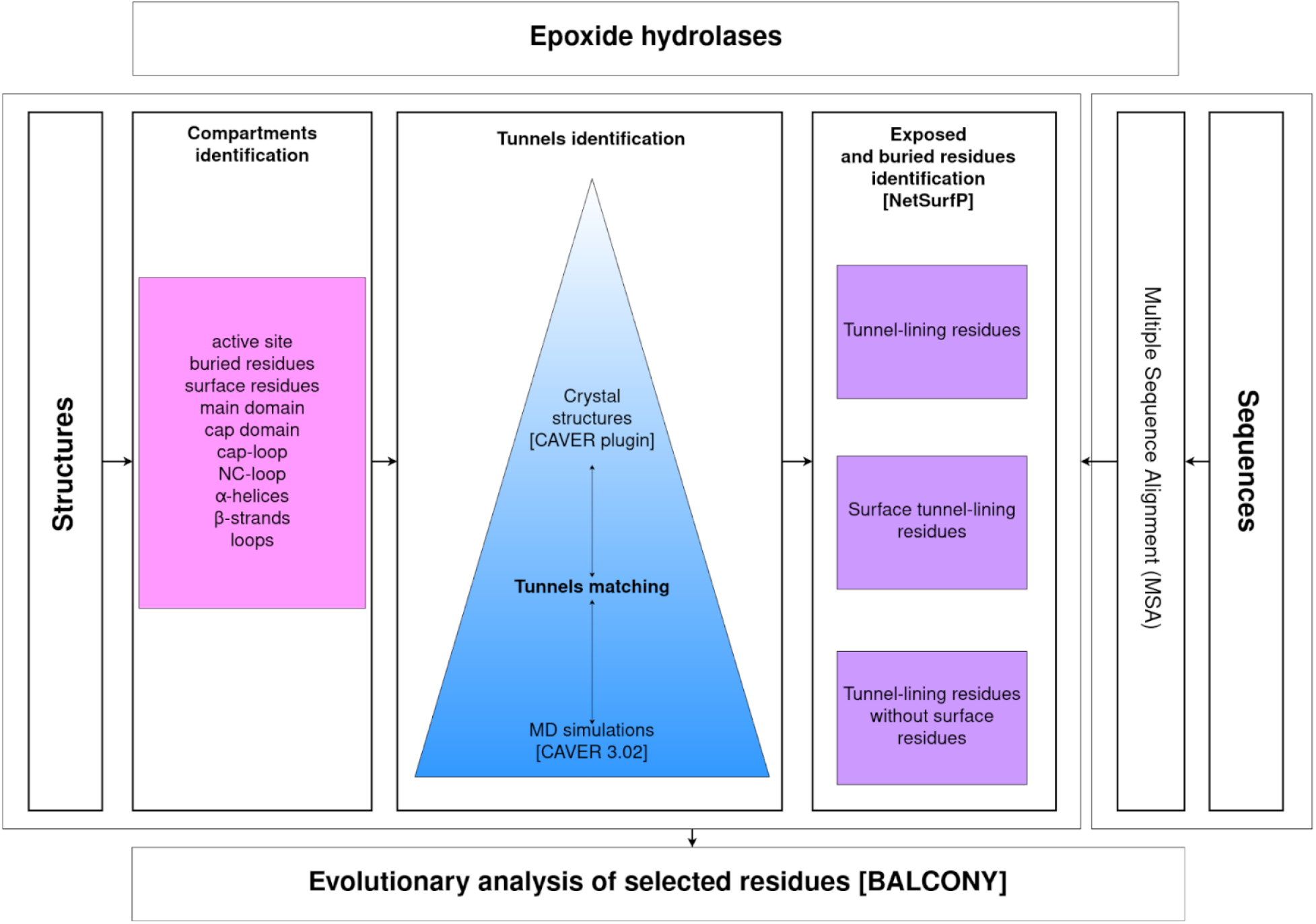
Research workflow.

### Obtaining protein structures for analysis

Seven unique and complete crystal structures were downloaded from the PDB database(39). The selected structures all belong to the α/β hydrolase superfamily, share the same core fold scheme(45), and consist of a main and a cap domains(40). Five structures represent different clades. They belong to clades of animals (*M. musculus* (msEH, PDB ID: 1CQZ)), *H. sapiens* (hsEH, PDB ID: 1S8O)), plants (*S. tuberosum* (StEH1, PDB ID: 2CJP)), fungi (*T. reesei* (TrEH, PDB ID: 5URO)), and bacteria (*B. megaterium* (bmEH, PDB ID: 4NZZ)). Two structures were collected from an unknown source organism in hot springs in Russia and China (Sibe-EH, PDB ID: 5NG7, CH65-EH, and PDB ID: 5NFQ).

### Structure preparation

Ligands were manually removed from each structure, and only one chain was used for the analysis. For the msEH and hsEH structures, only the C-terminal domain, with the hydrolytic activity, was used. Several referential structural compartments were selected for further analysis (see **Supplementary Table S2**): the active site; buried and surface residues; main and cap domains; cap-loop; NC-loop; and α-helices, β-strands, and loops. The definitions of the cap-loop and NC-loop were taken from the works of Barth *et al.*(40) and of Smit and Labuschagne(70). The NetSurfP service(44) was used to identify both buried and surface residues. Tunnels identified by CAVER software were also selected for further analysis.

### MD simulations

MD simulations for msEH (PDB ID: 1CQZ), hsEH (PDB ID: 1S8O), StEH1 (PDB ID: 2CJP), TrEH (PDB ID: 5URO), bmEH (PDB ID: 4NZZ), Sibe-EH (PDB ID: 5NG7), and CH65-EH (PDB ID: 5NFQ) were carried out according to the protocol described by Mitusińska *et al.*(47).

### CAVER analysis

Tunnel identification and analysis in each system were carried out using CAVER 3.02 software(46) in two steps: i) the crystal structure of the enzyme was analysed by the CAVER plugin for PyMOL(71); ii) tunnels were identified and analysed in 50,000 snapshots of multiple MD simulations by the standalone CAVER 3.02 software. The parameters used for both steps are shown in **Supplementary Table S14**. The tunnels found in MD simulations and in crystal structures were ranked and numbered based on their throughput value(46).

### Tunnels comparison

The tunnels identified during MD simulations and in crystal structures were compared based on the occurrence of tunnel-lining residues. For crystal structures, the occurrence was defined as the number of atoms of a particular amino acid that were identified as tunnel-forming atoms. For the sake of simplicity, no weighting scheme was used: Cα, backbone atoms, and side-chain atoms were considered to be of the same importance. For MD simulations’ results, tunnel occurrence was defined differently: as the number of MD snapshots in which particular amino acid was detected for a particular tunnel cluster. Therefore, this number could vary between 1 and the number of MD snapshots (50,000 in the performed analyses).

Despite the different definitions of occurrence used for crystal structures and for MD results, interpretation can be conducted in exactly the same way for each. Therefore, the above-defined occurrences can be directly used for fine-tuning the list of residues that form tunnels, i.e. by applying certain threshold values. In this study, the threshold was a number in the open range (0, 1) and amino acids were retained only if the condition

o > max(o) × τ

was satisfied, where o is the occurrence and τ is the threshold value.

For sets of tunnels detected in both the crystal structures and in the MD data, a distance matrix was calculated using the Jaccard distance formula(72):

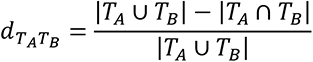

where T_A_ and T_B_ are A and B tunnels, respectively, and d is the Jaccard distance.

Elements of the matrix with lower distance values correspond to crystal structures/MD data pairs of similar tunnels. Further improvements in distance calculation accuracy were achieved by fine-tuning the tunnels’ amino acids with thresholds. For each of the compared pairs, two independent thresholds were used, and τ values for both lists of tunnel-forming residues in the crystal structure and MD simulations were scanned in the range of [0.05, 0.95] with a step of 0.05 (361 combinations in total). The combination of τ values which yielded the minimal distance was selected as the optimal one.

### Obtaining protein sequences, and MSA

Each of the amino acid sequences of the selected sEHs (PDB IDs: 1S8O, 1CQZ, 2CJP, 4NZZ, 5URO, 5NFQ, and 5NG7) was used as a separate query for a BLAST(73) search of similar protein sequences. The obtained results were merged and duplicates were removed, providing 1484 unique sequences (including those primarily selected). The 12 outlying sequences were detected and individually checked in the Uniprot database(74). Nine sequences were trimmed according to the hydrolase domain, and three were removed since there was no information or similarity with other sequences. Next, in order to eliminate proteins other than EHs from the set of sequences, the conserved motifs described by van Loo *et al.*(55) were used, and only sequences with motifs H-G-X-P and G-X-Sm-X-S/T were preserved (where X is usually an aromatic residue, and Sm is a small residue). As a result, 29 sequences were discarded during the analysis. In the last step of MSA preparation, additional domains (e.g. phosphatase domain) were removed. To detect sequences with an additional domain, a histogram of sequence lengths was prepared, and long sequences (> 420 residues) were trimmed all at once in a temporary MSA. In the end, a final MSA of 1455 epoxide hydrolase sequences was prepared with Clustal Omega(75) using default parameters (Supplementary Figure S11).

### BALCONY analysis

BALCONY (Better ALignment CONsensus analYsis)(41), an R package, was used to analyse the MSA and map selected structural compartments/tunnels onto the correct positions in aligned reference UniProt sequences. The Schneider metric(42) was calculated for each alignment position. Selected structures of *M. musculus*, *H. sapiens*, *S. tuberosum*, *T. reesei*, and *B. megaterium* sEHs, as well as the two thermophilic enzymes collected in hot springs (respective PDB IDs: 1CQZ, 1S8O, 2CJP, 5URO, 4NZZ, 5NG7, and 5NFQ), were divided into compartments/tunnels as shown in **Supplementary Table S1** and **Supplementary Tables S5–S11**. The compartment/tunnel residues were then appropriately mapped with MSA, and Schneider entropy values were collected for each position in the MSA.

### Variability analysis

To assess the variability of a particular tunnel/compartment, their positions were compared with selected positions of the MSA. The MSA was trimmed only to positions where at least one residue was present of the seven structures (PDB IDs: 1CQZ, 1S8O, 2CJP, 5URO, 4NZZ, 5NG7, and 5NFQ) (**Supplementary Figure S12**). The MSA containing 1455 sequences was trimmed from 722 to 419 positions. This way, for each comparison, Schneider entropy values of a compartment/tunnel positions were compared to the Schneider entropy values of selected positions of the MSA in which were present: i) neither one of residues of the currently analysed compartment/tunnel, and ii) at least one residue of the seven analysed structures. In order to determine whether a compartment was to be classed as variable, a median distance was calculated, which was defined as a difference between medians of Schneider entropy values of a selected compartment/tunnel and the selected positions in the MSA. If the median distance was > 0, then the analysed compartment was considered variable. To compare the distributions of entropy scores of analysed compartments/tunnels with the distribution of the selected positions of the MSA, the Epps–Singleton two-sample test(43) was used. The advantage of this test is the comparison of the empirical characteristic functions (the Fourier transform of the observed distribution function) instead of the observed distributions. The comparison analysis was performed using the es.test() function from GitHub repository(76). In an attempt to visualise the variability of selected tunnels (**Figure 5**), the collected entropy values of selected tunnel-lining residues without the surface residues and the selected MSA positions were sorted separately, and cumulative distribution functions (CDF) were calculated. For each position in the selected tunnel, a paired one from the selected position in the MSA was found, based on the minimal CDF. Plots of CDF as a function of entropy score were prepared.

**Supplementary Table S1.** List of amino acids forming particular compartments of analysed protein structures.

**Supplementary Table S2.** Differences in median Schneider entropy values between the median entropy values of the selected proteins’ compartments and remaining positions of the trimmed Multiple Sequence Alignment (MSA).

**Supplementary Table S3.** List of corresponding tunnels identified in both the crystal structure and in the MD simulation for each analysed protein structure.

**Supplementary Figure S1**. Issue related to identification of asymmetrical tunnels based on the example of the Tc/m tunnels identified by CAVER software during MD simulations.

**Supplementary Table S4.** Comparison of maximal bottleneck radii measured in corresponding tunnels identified in both the crystal structure (CR) and in the MD simulation (MD) for each protein structure.

**Supplementary Figure S2.** Correlation between maximal bottleneck radii measured in corresponding tunnels identified in both the crystal structure and in the MD simulation for each protein structure.

**Supplementary Table S5.** List of amino acids forming analysed tunnels in msEH structure.

**Supplementary Table S6.** List of amino acids forming analysed tunnels in hsEH structure.

**Supplementary Table S7.** List of amino acids forming analysed tunnels in StEH1 structure.

**Supplementary Table S8.** List of amino acids forming analysed tunnels in TrEH structure.

**Supplementary Table S9.** List of amino acids forming analysed tunnels in bmEH structure.

**Supplementary Table S10.** List of amino acids forming analysed tunnels in Sibe-EH structure.

**Supplementary Table S11.** List of amino acids forming analysed tunnels in CH65-EH structure.

**Supplementary Figure S3.** The distribution of the entropy values and the median entropy values of analysed tunnels and their parts and the remaining positions of the trimmed MSA for the *M. musculus* soluble epoxide hydrolase (msEH).

**Supplementary Figure S4.** The distribution of the entropy values and the median entropy values of analysed tunnels and their parts and the remaining positions of the trimmed MSA for the *H. sapiens* soluble epoxide hydrolase (hsEH).

**Supplementary Figure S5.** The distribution of the entropy values and the median entropy values of analysed tunnels and their parts and the remaining positions of the trimmed MSA for the *S. tuberosum* soluble epoxide hydrolase (StEH1).

**Supplementary Figure S6.** The distribution of the entropy values and the median entropy values of analysed tunnels and their parts and the remaining positions of the trimmed MSA for the *T. reesei* soluble epoxide hydrolase (TrEH).

**Supplementary Figure S7.** The distribution of the entropy values and the median entropy values of analysed tunnels and their parts and the remaining positions of the trimmed MSA for the *B. megaterium* soluble epoxide hydrolase (bmEH).

**Supplementary Figure S8.** The distribution of the entropy values and the median entropy values of analysed tunnels and their parts and the remaining positions of the trimmed MSA for the thermophilic enzyme collected in hot springs in Russia (Sibe-EH).

**Supplementary Figure S9.** The distribution of the entropy values and the median entropy values of analysed tunnels and their parts and the remaining positions of the trimmed MSA for the thermophilic enzyme collected in hot springs in China (CH65-EH).

**Supplementary Table S12.** Differences in Schneider entropy values between the median distance of selected tunnel-lining residues and the median distances of the remaining positions of the trimmed Multiple Sequence Alignment (MSA).

**Supplementary Table S13.** Entropy values for selected tunnels Tm1 from StEH1, Tc/m1 from hsEH, and Tc/m_back from bmEH.

**Supplementary Figure S10.** The open and closed position of the F497 residue of hsEH.

**Supplementary Table S14.** The list of parameters set for both CAVER plugin and CAVER 3.0 tunnels identification for each of the analysed systems.

**Supplementary Figure S11**. Representation of the created Multiple Sequence Alignment (MSA) of the epoxide hydrolases sequences.

**Supplementary Figure S12**. Representation of the trimmed Multiple Sequence Alignment (MSA) of the epoxide hydrolases sequences.

## Acknowledgement

This work was supported by the National Science Centre, Poland (grant number DEC-2013/10/E/NZ1/00649) (AG, MB, TM, KM).

## References

1. Kimura M, Ohta T. On Some Principles Governing Molecular Evolution. Proc Natl Acad Sci U S A. 1974;71(7):2848–52.

2. Tseng YY, Liang J. Estimation of Amino Acid Residue Substitution Rates at Local Spatial Regions and Application in Protein Function Inference: A Bayesian Monte Carlo Approach. Mol Biol Evol. 2005;23(2):421–36.

3. Franzosa EA, Xia Y. Structural determinants of protein evolution are context-sensitive at the residue level. Mol Biol Evol. 2009;26(10):2387–95.

4. Jäckel C, Kast P, Hilvert D. Protein Design by Directed Evolution. Annu Rev Biophys. 2008;37:153–73.

5. Damborsky J, Brezovsky J. Computational tools for designing and engineering enzymes. Curr Opin Chem Biol. 2014;19:8–16.

6. Garcia-Guevara F, Avelar M, Ayala M, Segovia L. Computational Tools Applied to Enzyme Design − a review. Biocatalysis. 2015;1:109–17.

7. Hochberg GKA, Thornton JW. Reconstructing Ancient Proteins to Understand the Causes of Structure and Function. Annu Rev Biophys. 2017;46(1):247–69.

8. Siddiq MA, Hochberg GK, Thornton JW. Evolution of protein specificity: insights from ancestral protein reconstruction. Curr Opin Struct Biol. 2017;47:113–22.

9. Arenas M, Bastolla U. ProtASR2: Ancestral reconstruction of protein sequences accounting for folding stability. Methods Ecol Evol. 2019;11(2):248–57.

10. Cuesta SM, Rahman SA, Furnham N, Thornton JM. The Classification and Evolution of Enzyme Function. Biophys J. 2015;109(6):1082–6.

11. Guney E, Tuncbag N, Keskin O, Gursoy A. HotSprint: database of computational hot spots in protein interfaces. Nucleic Acids Res. 2008;36(Database):D662–6.

12. Pavelka A, Chovancova E, Damborsky J. HotSpot Wizard: a web server for identification of hot spots in protein engineering. Nucleic Acids Res. 2009;37(Web Server):W376–83.

13. Verma R, Schwaneberg U, Roccatano D. MAP2.03D: A Sequence/Structure Based Server for Protein Engineering. ACS Synth Biol. 2012;1(4):139–50.

14. Martinez R, Schwaneberg U. A roadmap to directed enzyme evolution and screening systems for biotechnological applications. Biol Researcg. 2013;46(4).

15. Amaurys A, Bartlett IGJ, Hegedüs Z, Dutt S, Hobot F, Horner KA, et al. Predicting and Experimentally Validating Hot-Spot Residues at Protein–Protein Interfaces. ACS Chem Biol. 2019;14(10):2252–63.

16. Ramsey DC, Scherrer MP, Zhou T, Wilke CO. The Relationship Between Relative Solvent Accessibility and Evolutionary Rate in Protein Evolution. Genetics [Internet]. 2011 Jun;188(2):479–88. Available from: http://www.genetics.org/lookup/doi/10.1534/genetics.111.128025

17. Shahmoradi A, Sydykova DK, Spielman SJ, Jackson EL, Dawson ET, Meyer AG, et al. Predicting Evolutionary Site Variability from Structure in Viral Proteins: Buriedness, Packing, Flexibility, and Design. J Mol Evol [Internet]. 2014 Oct 13;79(3–4):130–42. Available from: http://link.springer.com/10.1007/s00239-014-9644-x

18. Sitbon E, Pietrokovski S. Occurrence of protein structure elements in conserved sequence regions. BMC Struct Biol. 2007;7(3):1–15.

19. Liu J, Tan H, Rost B. Loopy Proteins Appear Conserved in Evolution. J Mol Biol [Internet]. 2002 Sep;322(1):53–64. Available from: https://linkinghub.elsevier.com/retrieve/pii/S0022283602007362

20. Goldman N, Thorne JL, Jones DT. Assessing the Impact of Secondary Structureand Solvent Accessibility on Protein Evolution. Genetics. 1998;149:445–58.

21. Siltberg-Liberles J, Grahnen JA, Liberles DA, Siltberg-Liberles J, Grahnen JA, Liberles DA. The Evolution of Protein Structures and Structural Ensembles Under Functional Constraint. Genes (Basel) [Internet]. 2011 Oct 28 [cited 2018 Oct 24];2(4):748–62. Available from: http://www.mdpi.com/2073-4425/2/4/748

22. Jack BR, Meyer AG, Echave J, Wilke CO. Functional Sites Induce Long-Range Evolutionary Constraints in Enzymes. PLoS Biol [Internet]. 2016 [cited 2018 Oct 24];14(5):1002452. Available from: https://journals.plos.org/plosbiology/article/file?id=10.1371/journal.pbio.1002452&type=printable

23. Subramanian K, Mitusińska K, Raedts J, Almourfi F, Joosten H-J, Hendriks S, et al. Distant Non-Obvious Mutations Influence the Activity of a Hyperthermophilic Pyrococcus furiosus Phosphoglucose Isomerase. Biomolecules [Internet]. 2019 May 31;9(6):212. Available from: https://www.mdpi.com/2218-273X/9/6/212

24. Otten R, Liu L, Kenner LR, Clarkson MW, Mavor D, Tawfik DS, et al. Rescue of conformational dynamics in enzyme catalysis by directed evolution. Nat Commun. 2018 Dec;9(1):1314.

25. Petrović D, Risso VA, Kamerlin SCL, Sanchez-Ruiz JM. Conformational dynamics and enzyme evolution. J R Soc Interface. 2018 Jul;15(144):20180330.

26. Carlson GM, Fenton AW. What Mutagenesis Can and Cannot Reveal About Allostery. Biophys J. 2016 May;110(9):1912–23.

27. Weinkam P, Chen YC, Pons J, Sali A. Impact of Mutations on the Allosteric Conformational Equilibrium. J Mol Biol. 2013 Feb;425(3):647–61.

28. Marques SM, Daniel L, Buryska T, Prokop Z, Brezovsky J, Damborsky J. Enzyme Tunnels and Gates As Relevant Targets in Drug Design. Med Res Rev [Internet]. 2017 Sep;37(5):1095–139. Available from: http://doi.wiley.com/10.1002/med.21430

29. Kokkonen P, Bednar D, Pinto G, Prokop Z, Damborsky J. Engineering enzyme access tunnels. Biotechnol Adv [Internet]. 2019 Nov;37(6):107386. Available from: https://linkinghub.elsevier.com/retrieve/pii/S0734975019300679

30. Kingsley LJ, Lill MA. Substrate tunnels in enzymes: Structure-function relationships and computational methodology. Proteins Struct Funct Bioinforma [Internet]. 2015 Apr;83(4):599–611. Available from: http://doi.wiley.com/10.1002/prot.24772

31. Nakamura A, Yao M, Chimnaronk S, Sakai N, Tanaka I. Ammonia Channel Couples Glutaminase with Transamidase Reactions in GatCAB. Science (80-) [Internet]. 2006 Jun 30;312(5782):1954–8. Available from: https://www.sciencemag.org/lookup/doi/10.1126/science.1127156

32. Kim J, Raushel FM. Perforation of the Tunnel Wall in Carbamoyl Phosphate Synthetase Derails the Passage of Ammonia between Sequential Active Sites. Biochemistry. 2004;43:5334–40.

33. Thangapandian S, John S, Lee Y, Arulalapperumal V, Lee KW. Molecular Modeling Study on Tunnel Behavior in Different Histone Deacetylase Isoforms. Gaetano C, editor. PLoS One [Internet]. 2012 Nov 29;7(11):e49327. Available from: https://dx.plos.org/10.1371/journal.pone.0049327

34. Zawaira A, Coulson L, Gallotta M, Karimanzira O, Blackburn J. On the deduction and analysis of singlet and two-state gating-models from the static structures of mammalian CYP450. J Struct Biol [Internet]. 2011 Feb;173(2):282–93. Available from: https://linkinghub.elsevier.com/retrieve/pii/S1047847710002959

35. Nardini M, Dijkstra BW. α/β Hydrolase fold enzymes: the family keeps growing. Curr Opin Struct Biol [Internet]. 1999 Dec;9(6):732–7. Available from: https://linkinghub.elsevier.com/retrieve/pii/S0959440X99000378

36. Marchot P, Chatonnet A. Enzymatic Activity and Protein Interactions in Alpha/Beta Hydrolase Fold Proteins: Moonlighting Versus Promiscuity. Protein Pept Lett [Internet]. 2012 Feb 1;19(2):132–43. Available from: http://www.eurekaselect.com/openurl/content.php?genre=article&issn=0929-8665&volume=19&issue=2&spage=132

37. Bauer TL, Buchholz PCF, Pleiss J. The modular structure of α/β-hydrolases. FEBS J [Internet]. 2020 Mar 10;287(5):1035–53. Available from: https://onlinelibrary.wiley.com/doi/abs/10.1111/febs.15071

38. Holmquist M. Alpha Beta-Hydrolase Fold Enzymes Structures, Functions and Mechanisms. Curr Protein Pept Sci [Internet]. 2000 Sep 1;1(2):209–35. Available from: http://www.ingentaselect.com/rpsv/cgi-bin/cgi?ini=xref&body=linker&reqdoi=10.2174/1389203003381405

39. Berman HM. The Protein Data Bank. Nucleic Acids Res [Internet]. 2000 Jan 1;28(1):235–42. Available from: https://academic.oup.com/nar/article-lookup/doi/10.1093/nar/28.1.235

40. Barth S, Fischer M, Schmid RD, Pleiss J. Sequence and structure of epoxide hydrolases: A systematic analysis. Proteins Struct Funct Bioinforma [Internet]. 2004 Apr 2;55(4):846–55. Available from: http://doi.wiley.com/10.1002/prot.20013

41. Płuciennik A, Stolarczyk M, Bzówka M, Raczyńska A, Magdziarz T, Góra A. BALCONY: an R package for MSA and functional compartments of protein variability analysis. BMC Bioinformatics [Internet]. 2018 Dec 14;19(1):300. Available from: https://bmcbioinformatics.biomedcentral.com/articles/10.1186/s12859-018-2294-z

42. Sander C, Schneider R. Database of homology-derived protein structures and the structural meaning of sequence alignment. Proteins Struct Funct Genet [Internet]. 1991 Jan;9(1):56–68. Available from: http://doi.wiley.com/10.1002/prot.340090107

43. Epps TW, Singleton KJ. An omnibus test for the two-sample problem using the empirical characteristic function. J Stat Comput Simul [Internet]. 1986 Dec;26(3–4):177–203. Available from: http://www.tandfonline.com/doi/abs/10.1080/00949658608810963

44. Klausen MS, Jespersen MC, Nielsen H, Jensen KK, Jurtz VI, Sønderby CK, et al. NetSurfP-2.0: Improved prediction of protein structural features by integrated deep learning. Proteins Struct Funct Bioinforma [Internet]. 2019 Jun 9;87(6):520–7. Available from: https://onlinelibrary.wiley.com/doi/abs/10.1002/prot.25674

45. Ollis DL, Cheah E, Cygler M, Dijkstra B, Frolow F, Franken SM, et al. The α / β hydrolase fold. “Protein Eng Des Sel [Internet]. 1992;5(3):197–211. Available from: https://academic.oup.com/peds/article-lookup/doi/10.1093/protein/5.3.197

46. Chovancova E, Pavelka A, Benes P, Strnad O, Brezovsky J, Kozlikova B, et al. CAVER 3.0: A Tool for the Analysis of Transport Pathways in Dynamic Protein Structures. Prlic A, editor. PLoS Comput Biol [Internet]. 2012 Oct 18;8(10):e1002708. Available from: https://dx.plos.org/10.1371/journal.pcbi.1002708

47. Mitusińska K, Wojsa P, Bzówka M, Raczyńska A, Bagrowska W, Samol A, et al. Structure-function relationship between soluble epoxide hydrolases structure and their tunnel network. Comput Struct Biotechnol J [Internet]. 2022;20:193–205. Available from: https://linkinghub.elsevier.com/retrieve/pii/S2001037021005225

48. Bzówka M, Mitusińska K, Hopko K, Góra A. Computational insights into the known inhibitors of human soluble epoxide hydrolase. Drug Discov Today [Internet]. 2021 Aug;26(8):1914–21. Available from: https://linkinghub.elsevier.com/retrieve/pii/S135964462100252X

49. Mitusińska K, Magdziarz T, Bzówka M, Stańczak A, Gora A. Exploring Solanum tuberosum Epoxide Hydrolase Internal Architecture by Water Molecules Tracking. Biomolecules [Internet]. 2018 Nov 12;8(4):143. Available from: http://www.mdpi.com/2218-273X/8/4/143

50. Kong X-D, Yuan S, Li L, Chen S, Xu J-H, Zhou J. Engineering of an epoxide hydrolase for efficient bioresolution of bulky pharmaco substrates. Proc Natl Acad Sci [Internet]. 2014 Nov 4;111(44):15717–22. Available from: http://www.pnas.org/cgi/doi/10.1073/pnas.1404915111

51. Caetano-Anollés G, Wang M, Caetano-Anollés D, Mittenthal JE. The origin, evolution and structure of the protein world. Biochem J [Internet]. 2009 Feb 1;417(3):621–37. Available from: https://portlandpress.com/biochemj/article/417/3/621/44492/The-origin-evolution-and-structure-of-the-protein

52. Minary P, Levitt M. Probing Protein Fold Space with a Simplified Model. J Mol Biol [Internet]. 2008 Jan;375(4):920–33. Available from: https://linkinghub.elsevier.com/retrieve/pii/S0022283607014581

53. Suplatov DA, Besenmatter W, Svedas VK, Svendsen A. Bioinformatic analysis of alpha/beta-hydrolase fold enzymes reveals subfamily-specific positions responsible for discrimination of amidase and lipase activities. Protein Eng Des Sel [Internet]. 2012 Nov 1;25(11):689–97. Available from: https://academic.oup.com/peds/article-lookup/doi/10.1093/protein/gzs068

54. Heikinheimo P, Goldman A, Jeffries C, Ollis DL. Of barn owls and bankers: a lush variety of α/β hydrolases. Structure [Internet]. 1999 Jun;7(6):R141–6. Available from: https://linkinghub.elsevier.com/retrieve/pii/S0969212699800793

55. van Loo B, Kingma J, Arand M, Wubbolts MG, Janssen DB. Diversity and Biocatalytic Potential of Epoxide Hydrolases Identified by Genome Analysis. Appl Environ Microbiol [Internet]. 2006 Apr 1;72(4):2905–17. Available from: http://aem.asm.org/cgi/doi/10.1128/AEM.72.4.2905-2917.2006

56. Dimitriou PS, Denesyuk A, Takahashi S, Yamashita S, Johnson MS, Nakayama T, et al. Alpha/beta-hydrolases: A unique structural motif coordinates catalytic acid residue in 40 protein fold families. Proteins Struct Funct Bioinforma [Internet]. 2017 Oct;85(10):1845–55. Available from: http://doi.wiley.com/10.1002/prot.25338

57. Lindberg D, Ahmad S, Widersten M. Mutations in salt-bridging residues at the interface of the core and lid domains of epoxide hydrolase StEH1 affect regioselectivity, protein stability and hysteresis. Arch Biochem Biophys [Internet]. 2010 Mar;495(2):165–73. Available from: https://linkinghub.elsevier.com/retrieve/pii/S0003986110000202

58. Jeon J, Nam H-J, Choi YS, Yang J-S, Hwang J, Kim S. Molecular Evolution of Protein Conformational Changes Revealed by a Network of Evolutionarily Coupled Residues. Mol Biol Evol [Internet]. 2011 Sep;28(9):2675–85. Available from: https://academic.oup.com/mbe/article-lookup/doi/10.1093/molbev/msr094

59. Schiøtt B, Bruice TC. Reaction Mechanism of Soluble Epoxide Hydrolase: Insights from Molecular Dynamics Simulations §. J Am Chem Soc [Internet]. 2002 Dec;124(49):14558–70. Available from: https://pubs.acs.org/doi/10.1021/ja021021r

60. Bahl CD, Morisseau C, Bomberger JM, Stanton BA, Hammock BD, O’Toole GA, et al. Crystal Structure of the Cystic Fibrosis Transmembrane Conductance Regulator Inhibitory Factor Cif Reveals Novel Active-Site Features of an Epoxide Hydrolase Virulence Factor. J Bacteriol [Internet]. 2010 Apr 1;192(7):1785–95. Available from: https://jb.asm.org/content/192/7/1785

61. Lindberg D, de la Fuente Revenga M, Widersten M. Temperature and pH Dependence of Enzyme-Catalyzed Hydrolysis of trans -Methylstyrene Oxide. A Unifying Kinetic Model for Observed Hysteresis, Cooperativity, and Regioselectivity. Biochemistry [Internet]. 2010 Mar 16;49(10):2297–304. Available from: https://pubs.acs.org/doi/10.1021/bi902157b

62. Hvorecny KL, Bahl CD, Kitamura S, Lee KSS, Hammock BD, Morisseau C, et al. Active-Site Flexibility and Substrate Specificity in a Bacterial Virulence Factor: Crystallographic Snapshots of an Epoxide Hydrolase. Structure [Internet]. 2017 May;25(5):697–707.e4. Available from: https://linkinghub.elsevier.com/retrieve/pii/S096921261730062X

63. Mowbray SL, Elfström LT, Ahlgren KM, Andersson CE, Widersten M. X-ray structure of potato epoxide hydrolase sheds light on substrate specificity in plant enzymes. Protein Sci [Internet]. 2006 Jul;15(7):1628–37. Available from: http://doi.wiley.com/10.1110/ps.051792106

64. Hasan K, Gora A, Brezovsky J, Chaloupkova R, Moskalikova H, Fortova A, et al. The effect of a unique halide-stabilizing residue on the catalytic properties of haloalkane dehalogenase DatA from Agrobacterium tumefaciens C58. FEBS J [Internet]. 2013 Jul;280(13):3149–59. Available from: http://doi.wiley.com/10.1111/febs.12238

65. Swint-Kruse L. Using Evolution to Guide Protein Engineering: The Devil IS in the Details. Biophys J [Internet]. 2016 Jul;111(1):10–8. Available from: https://linkinghub.elsevier.com/retrieve/pii/S0006349516303551

66. Chaloupková R, Sýkorová J, Prokop Z, Jesenská A, Monincová M, Pavlová M, et al. Modification of Activity and Specificity of Haloalkane Dehalogenase from Sphingomonas paucimobilis UT26 by Engineering of Its Entrance Tunnel. J Biol Chem [Internet]. 2003 Dec;278(52):52622–8. Available from: https://linkinghub.elsevier.com/retrieve/pii/S0021925820752749

67. Janfalk Carlsson Å, Bauer P, Dobritzsch D, Nilsson M, Kamerlin SCL, Widersten M. Laboratory-Evolved Enzymes Provide Snapshots of the Development of Enantioconvergence in Enzyme-Catalyzed Epoxide Hydrolysis. ChemBioChem [Internet]. 2016 Sep 15;17(18):1693–7. Available from: http://doi.wiley.com/10.1002/cbic.201600330

68. Aharoni A, Gaidukov L, Khersonsky O, Gould SM, Roodveldt C, Tawfik DS. The “evolvability” of promiscuous protein functions. Nat Genet [Internet]. 2005 Jan 28;37(1):73–6. Available from: http://www.nature.com/articles/ng1482

69. Brezovsky J, Babkova P, Degtjarik O, Fortova A, Gora A, Iermak I, et al. Engineering a de Novo Transport Tunnel. ACS Catal [Internet]. 2016 Nov 4;6(11):7597–610. Available from: https://pubs.acs.org/doi/10.1021/acscatal.6b02081

70. Smit M, Labuschagne M. Diversity of Epoxide Hydrolase Biocatalysts. Curr Org Chem [Internet]. 2006 Jul 1;10(10):1145–61. Available from: http://www.eurekaselect.com/openurl/content.php?genre=article&issn=1385-2728&volume=10&issue=10&spage=1145

71. Schrödinger. The PyMOL Molecular Graphic Systems. Schrödinger, LLC;

72. Gilbert G. Distance between Sets. Nature [Internet]. 1972 Sep;239(5368):174–174. Available from: http://www.nature.com/articles/239174c0

73. Altschul SF, Gish W, Miller W, Myers EW, Lipman DJ. Basic local alignment search tool. J Mol Biol [Internet]. 1990 Oct;215(3):403–10. Available from: https://linkinghub.elsevier.com/retrieve/pii/S0022283605803602

74. UniProt Consortium T. UniProt: the universal protein knowledgebase. Nucleic Acids Res [Internet]. 2018 Mar 16;46(5):2699–2699. Available from: https://academic.oup.com/nar/article/46/5/2699/4841658

75. Sievers F, Higgins DG. Clustal Omega, Accurate Alignment of Very Large Numbers of Sequences. In 2014. p. 105–16. Available from: http://link.springer.com/10.1007/978-1-62703-646-7_6

76. Ileppane. https://github.com/ileppane/statistics/blob/master/EStest.R. 2014.

