## Supplementary File for "Evolution of tunnels in α/β-hydrolase fold proteins – what can we learn from studying epoxide hydrolases?"

### Supplementary Information

**Supplementary Table S1**. List of amino acids forming particular compartments of analysed protein structures.

**Supplementary Table S2**. Differences in median Schneider entropy values between the median entropy values of the selected proteins' compartments and remaining positions of the trimmed Multiple Sequence Alignment (MSA).

**Supplementary Table S3**. List of corresponding tunnels identified in both the crystal structure and in the MD simulation for each analysed protein structure.

**Supplementary Figure S1**. Issue related to identification of asymmetrical tunnels based on the example of the Tc/m tunnels identified by CAVER software during MD simulations.

**Supplementary Table S4.** Comparison of maximal bottleneck radii measured in corresponding tunnels identified in both the crystal structure (CR) and in the MD simulation (MD) for each protein structure.

**Supplementary Figure S2.** Correlation between maximal bottleneck radii measured in corresponding tunnels identified in both the crystal structure and in the MD simulation for each protein structure.

**Supplementary Table S5.** List of amino acids forming analysed tunnels in msEH structure.

**Supplementary Table S6.** List of amino acids forming analysed tunnels in hsEH structure.

**Supplementary Table S7.** List of amino acids forming analysed tunnels in StEH1 structure.

**Supplementary Table S8.** List of amino acids forming analysed tunnels in TrEH structure.

**Supplementary Table S9.** List of amino acids forming analysed tunnels in bmEH structure.

**Supplementary Table S10.** List of amino acids forming analysed tunnels in Sibe-EH structure.

**Supplementary Table S11.** List of amino acids forming analysed tunnels in CH65-EH structure.

**Supplementary Figure S3.** The distribution of the entropy values and the median entropy values of analysed tunnels and their parts and the remaining positions of the trimmed MSA for the *M. musculus* soluble epoxide hydrolase (msEH).

**Supplementary Figure S4.** The distribution of the entropy values and the median entropy values of analysed tunnels and their parts and the remaining positions of the trimmed MSA for the *H. sapiens* soluble epoxide hydrolase (hsEH).

**Supplementary Figure S5.** The distribution of the entropy values and the median entropy values of analysed tunnels and their parts and the remaining positions of the trimmed MSA for the *S. tuberosum* soluble epoxide hydrolase (StEH1).

**Supplementary Figure S6.** The distribution of the entropy values and the median entropy values of analysed tunnels and their parts and the remaining positions of the trimmed MSA for the *T. reesei* soluble epoxide hydrolase (TrEH).

**Supplementary Figure S7.** The distribution of the entropy values and the median entropy values of analysed tunnels and their parts and the remaining positions of the trimmed MSA for the *B. megaterium* soluble epoxide hydrolase (bmEH).

**Supplementary Figure S8.** The distribution of the entropy values and the median entropy values of analysed tunnels and their parts and the remaining positions of the trimmed MSA for the thermophilic enzyme collected in hot springs in Russia (Sibe-EH).

**Supplementary Figure S9.** The distribution of the entropy values and the median entropy values of analysed tunnels and their parts and the remaining positions of the trimmed MSA for the thermophilic enzyme collected in hot springs in China (CH65-EH).

**Supplementary Table S12.** Differences in Schneider entropy values between the median distance of selected tunnel-lining residues and the median distances of the remaining positions of the trimmed Multiple Sequence Alignment (MSA).

**Supplementary Table S13.** Entropy values for selected tunnels Tm1 from StEH1, Tc/m1 from hsEH, and Tc/m_back from bmEH.

**Supplementary Figure S10.** The open and closed position of the F497 residue of hsEH.

**Supplementary Table S14.** The list of parameters set for both CAVER plugin and CAVER 3.0 tunnels identification for each of the analysed systems.

**Supplementary Figure S11**. Representation of the created Multiple Sequence Alignment (MSA) of the epoxide hydrolases sequences.

**Supplementary Figure S12**. Representation of the trimmed Multiple Sequence Alignment (MSA) of the epoxide hydrolases sequences.

**Supplementary Table S1**. List of amino acids forming particular compartments of analysed protein structures.

| Protein | Compartment | Residues |
|---|---|---|
| msEH - 1CQZ | Active site | D333, Y381, Y465, D495, H523 |
| | Buried amino acids | V240, V242, I246, l248-M253, G256, A258-W272, Q275-I276, A278-A280, F284-G295, S297, P300, Y306, M308, L310-C312, E314-V316, F318-L319, L322, I324, A327-L344, Y346, R349-V350, A352-P363, D364, V367, I373, I376, F389-F385, V390-A391, E394-L395, M399, T402-F403, F406-F407, S410, V418, A421, I424, I427-L428, T431-P432, L437, I441-T441, E445-I446, F448-I450, Q452-F453, F458, G460-T468, W472, C476, G480, I483, V485-A492, D495, V497-L498, P500, M502-S503, M506, W509-I510, L513, R515, I518, C521-P530, V533-N534, I536-I538, W540-L541 |
| | Surface amino acids | H237-Y239; T241; K243-G245; R247; G254-S255; P257; R273-Y274; P277; Q281-G283; D296; S298-S299; P301-E305; A307; E309; K313; T317; D320-K321; G323; P325-Q326; F345; P347-E348; R351; P353; P365-D366; S368-V372; R274-S375; P377-V378; Q386-F389; E392-A393; E396-N398; S400-R401; K404-S405; R408-A409; D411-A417; H419-K420; T422-E423; G425-G426; V429-N430; E433-N436; S438-K439; T442-E444; E447; Q451; K454-G547; R459; R466; E469; R470-N471; K473-S475; K477-L479; R481-K482; L484; E493-K494; I496; R499; E501; K504-N505; E507-K508; P511-F512; K514; G516-H517; E519-D520; T531-E532; Q535; K539; Q542-I554 |
| | Main domain | V235-S368; R470-I554 |
| | Cap domain | P369-Q469 |
| | Cap-loop | F407-T442 |
| | NC-loop | T346-S368 |
| | Alpha-helices | Y269-A282; I303-L322;<br>Y334-F345; P369-R374; N380-Q386; A391-E396; M399-F406; A421-I424; E443-K455; F458-G480;<br>P500-Y509; T525-Q542 |
| | Beta-sheets | S236-K243; I246-M253; A258-C262; R285-I289; V328-G331; V353-L356; A487-A492; K514-I518 |
| | Loops | V235; P244-G245; G254-P257; H263-S268; G283-F284; D290-E302; A307; G323-D333; T346-A352; N357-S368; S375-F379; E387-V390; K397-N398; F407-L420; G425-T442; T456-G457; R481-P486; E493-R499; I510-L513; E519-Y524; K529; T543-E544 |
| hsEH-1S8O | Active site | D335, Y383, Y466, D496, H524 |
| | Buried amino acids | V242, V244, V248, L250-L255, G258, A261-R275, Q277-I278, A280-A282, Y286-G297, S299, P302, Y308, M310, V312-C314, E316-V318, F320-L321, L324, L326, A329-L346, Y348, R351-V352, A354-P364, N366, M369, L372, I375, N378, F381-F387, V392--A393, E396-L397, L401, T404-F405, L408-F409, V416, M419, V422-C423, A425, L428-F429, S432-P433, L438, M441-T443, E446-I447, F449-V451, Q453-F454, F459, G461-M469, W473, C477, G481, I484, I486-A493, D496, V498-P501, M503-S504, M507, W510-I511, L514, R516-G517, I519, C522-Q527, D529-P531, V534-N535, I537-I539, W541-L542 |
| | Surface amino acids | S238-Y241; T243; K245-R247; R249; G256-S257; P259; Y276; P279; Q283-G285; E298; S300-A301; P303-E307; C309; E311; K315; T319; D322-K323; G325; S327- |

| | | |
|---|---|---|
| | | Q328; F347; P349-E350; R353; A365; P367-N368; S370-P371; E373-S374; K376-A377; P379-V380; Q388-G391; E394-A395; E398-N400; S402-R403; K406-S407; R410-S415; L417-S418; H420-K421; E424; G426-G427; V430-N431; E434-S437; S439-R440; E444-E445; Q448; Q452; K455-G458; R460; E470-N472; K474-A476; K478-L480; R482-K483; L485; E494-K495; F497; Q502; Q505-H506; E508-D509; P512-H513; K515; H518; E520-D521; M528; T532-E533; Q536; K540; D543-M555 |
| | Main domain | M235-P369; E469-M555 |
| | Cap domain | S370-N468 |
| | Cap-loop | F409-T443 |
| | NC-loop | T348-S370 |
| | Alpha-helices | Y271-A284; I305-L324;<br>Y336-F347; P349-R351;<br>P371-A377; P379-Q388; V392-Q399; L401-L408; V422-E424; E444-S457;<br>F459-K478; P501-Y510; P531-D543 |
| | Beta-sheets | H240-K245; V248-L255; A260-H265; R287-M291; V330-L358; A488-T492; L514-I519 |
| | Loops | H239; P246-R247; G256-P259; G266-S270; G285-Y286; D292-E304; C309; G325-A329; D335; Y348; V352-A354; N359-S370; N378; E389-G391; N400; F409-K421; G426-T443; G458; S479-P487; A493-V500; I511-H513; E520-K530; S544-N548 |
| StEH1 - 2CJP | Active site | D105, Y154, Y235, D265, H300 |
| | Buried amino acids | I4, V9, V11, L14, M16-A19, L21, G24, T26-W40, Q43-M44, Y46-A48, Y52-G63, T65, A68, F76, I78, L81-V82, D84-V86, L88-L89, I92, P94, B99-L116, R118, K121-S133, N136, V141, L145, Y149, H153-F158, V160-P161, I164, F168, A173, V176-L177, K179-L181, F191, G196-L197, I200-P201, W210-L211, E215-L216, Y218-A220, K222-F223, F228, G230-L238, W242, V253, V255-G262, D265, V267-Y268, I270, G272, A273, Y276-I277, F282, V286-P287, E290, V292-L295, A298-P307, I310-S311, H313-I314, F317-I318 |
| | Surface amino acids | K3; E5-M8; A10; N12-N15; E20; G22-E23; P25; R41-H42; V45; E49-G51; D64; T66-G67; P69; L70-K75; S77; L79-H80; G83; A87; E90-A91; A93; N95-K98; F117; P119-D120; K134-R135; P137-N140; V142-G144; K146-I148; G150-D152; Q159; G162-E163; E165-E167; A169-G172; K174-S175; K178; T182-Y190; P192-K195; E198-A199; D202-S209; S212-E214; D217; N221; E224-G227; T229; P239-N241; E243-Q252; K254; E263-F264; L266; H269; P271; K274-E275; H278-G281; K283-D285; L289; E291; E296-G297; H308-E309; K312; Y315-D316; Q319-F321 |
| | Main domain | K2-N140; L238-F321 |
| | Cap domain | V141-A237 |
| | Cap-loop | T128-S212 |
| | NC-loop | R118-N140 |
| | Alpha-helices | W37-R50; P73-I92; W106-K121; V141-Y149; Y154-Q159; E163-I171; A173-T182; L197-A199; V205-L207; E213T226; F228-T245; A273-N279; G281-D285; V302-K320 |

| | | |
|---|---|---|
| | Beta-sheets | E5-V11; L14-L21; T26-H31; R53-P57; F100-H104; L125-L128; T257-G262; V293-L295 |
| | Loops | K3-I4; N12-G13; G22-P25; G32-L36; G51-Y52; D58-D72; S77; A93-V99; D105; R118; V122-A124; S129-N140; G150-H153; V160-G162; G172; Y183-G196; I200-P204; S208-S212; G227; A246-P256; E263-G272; G280; V286-V292; E296-F301; R306; F321 |
| TrEH - 5URO | Active site | D1016 Y167, Y252, D286, H313 |
| | Buried amino acids | L6, V13, Y15-E16, K18, I20, K23, Y25-E31, L37-W53, Q56-I57, Y59-M61, F65-A76, T78, P81, F87, L89, S91-A100, F103, G105, G108, I110-Y127, H129, L132-P145, Y150, L153, I156, M162-L171, G173, V176-E177, I180-Q181, M185-L186, F189-F190, A192-F194, G198, G201, F205, V211-H212, V215-L216, I219, L224-L225, E229-L230, Y232-V234, Q236-Y237, E244-E256, A259, M263, K267, P271, F274, M276-A283, D286, A288-P291, M293-S294, G296-M297, F300-Y301, L304, R306-V309, A311-G320, N324, V326-F328, W330-L331 |
| | Surface amino acids | M1-K5; K7-R12; K14; T17; Q19; R21-G22; T24; P32-K36; R54-H55; P58; S62-G64; G77; D79-A80; R82-Q86; T88; K90; A94; R101-S102; V104; Q106-D107; Q109; Y128; P130-E131; L146-E149; K151-P152; E154-D155; V157-H161; K172; P174-D175; A178-R179; G182-D184; R187-R188; R191; G195-R197; P199-N200; E202-G204; S206-G210; F213-D214; D217-K218; G220-P223; D226-Q228; E231; E235; A238-P243; L257-N258; K2601-E262; D264-A66; N268-P270; L272-R273; E275; S284-K285; N287; A292; K295; D298-A299; K302-D303; T305; D310; A319; D321-E322; R325; E329; N332-G336 |
| | Main domain | M1-L153; T254-G336 |
| | Cap domain | E154-R253 |
| | Cap-loop | G195-D226 |
| | NC-loop | H129-K151 |
| | Alpha-helices | A50-L63; L84-F87; L89-V104; W117-Y128; L153-A159; L163-K172; D175-R179; K183-F194; F213-K218; E227-L239; E244-K267; P291-S294; M297-F300; A315-A334 |
| | Beta-sheets | K14-I20; K23-G30; T39-H44; Q66-P70; I110-H115; A135-V139; A278-A283; T305-V309 |
| | Loops | M1-V13; R21-G22; E31-G38; G45-M49; G64-F65; N71-D83; T88; G105-Q109; D116; H129-K134; C140-P152; G160-M162; G173-P174; I180-G182; G195-H212; I219-D226; Q240-P243; N268-P277; S284-P290; K295-H296; Y301-L304; D310-W314; W335-G336 |
| bmEH - 4NZZ | Active site | D97, Y144, Y203, D239, H267 |
| | Buried amino acids | I6, V8, V11, L13-S17, G21, L23-R38, Q40-I41, F44-S45, F48-N59, S61, P64, Y70, I72, V74-V76, D78-R80, V82-I83, L86, Y88, C91-Y108, Y110-P111, Y113-P125, F128, L132, K136, Q139-M145, W147-F148, V153-Q154, Y156-M157, F162, G164-R166, L168-V169, P172-V174, L179, D183-V184, A186-M188, W191, S195-V196, S198-K207, F209, L214, F220, E224, L227, I229-G236, D239, T241-F242, P244, N246-L247, I250, Y253-V254, I257, V259-H260, L262, A265-P274, V277-N278, V280-M281, F284-L285 |
| | Surface amino acids | M1-Y5; N7; N9-G10; N12; K18-Q20; E22; H39; D42-E43; N46-D47; L60; E62-K63; S65-S69; E71; D73; E77; Q81; E84-G85; G87; S89-S90; R109; E112; Y126-T127; M129-E131; R133-N135; N137-Q138; K146; Q149-E152; D155; E158-N161; S163; |

| | | |
|---|---|---|
| | | K167; I170-D171; K175-Y178; T180-D182; Q185; N189-S190; E192-G194; L197; I208; T210-D213; R215-L219; P221; L222-E223; E225-V226; N228; N237-Q238; P240; M243; E245; D248-G249; E251-E252; P255-N256; S258; R261; A263-E264; Q275-E276; N279; W282-N283; N286-K287 |
| | Main domain | M1-N135; K207-K287 |
| | Cap domain | K136-L206 |
| | Cap-loop | K176-A181 |
| | NC-loop | T110-L132 |
| | Alpha-helices | S34-S45; I72-L86; W98- Y113; Y126-Q149; Q151-E158; S164-L168; I170-K176; A181-N193; V196-N205; E211-R215; P244-L247; O250-Y253; P274-N286 |
| | Beta-sheets | S2-V8; V11-K18; L23-H28; H49-D54; T92-H96; K116-A119; V231-L262 |
| | Loops | M1; N9-G10; G19-E22; G29-F33; N46-F48; L55-E71; G87-C91; D97; Y110; V114-Q115; F120-P125; N135; K150; R159-F162; V169; G177-T180; G194-S195; L206-T210; R216-P230; N237-M243; D248-G249; V254-N256; A263-K273; K287 |
| Sibe-EH-5NG7 | Active site | D101, Y148, Y209, D241, H270 |
| | Buried amino acids | L5, E8, V10, V12, I15, M17-V20, G25, L27-R42, Q44-I45, L48-A49, F52-N63, T65, P68, Y74, L76, L79-A80, D82-L84, L86-I87, L90, E92, A95-A112, N114, A117-V118, K120-P129, T136, S139-L140, L143-Q144, S146-F152, I157-P158, K160-L162, F169-L170, M173-L174, S177-F178, L184, D188-L189, Y192-V193, A195-W196, A201-L202, S204-L213, F219, K222, F226, I229, V231-G238, D241, A243-I244, K246, L248-I249, M252-E253, I256, Y260, I262-F265, C268-Q273, E275-P277, V280-R281, H283-E285, F287-I288 |
| | Surface amino acids | N2-M4; K6-H7; Y9; K11; N13-G14; K16; T21-K24; K26; F43; P46-A47; K50-H51; E64; D66-K67; E69-N73; R75; D77-L78; K81; G85; K88-A89; G91; E93-H94; F113; P115-Q116; E119; K130-R135; K137-N138; R141-Q142; K145; Q153-N156; E159; S163-A168; K171-N172; I175-Q176; V179-L183; T185-E187; R190-I191; D194; S197-G200; T203; N214-I218; S220-E221; T223-V225; P227-K228; K230; E239-K240; V242; S245; D247; V250-N251; D254-F255; E257-P250; S261; P266-E267; L274; E278-L279; K282; E286; L289-I293 |
| | Main domain | M1-P127; E221-I293 |
| | Cap domain | H128-S220 |
| | Cap-loop | F178-T185 |
| | NC-loop | N114-P127 |
| | Alpha-helices | W38-A49; L76-L90; W102-A117; P129-Q153; I157-S163; A168-S177; E186-S197; A201-I218; K246-I249; M252-F255; P277-L289 |
| | Beta-sheets | K6-V12; I15-Q22; L27-H32; R53-P57; V96-H100; K120-L124; T233-G238; Y260-F265 |

| | | |
|---|---|---|
| | Loops | N2-L5; N13-G14; G23-K26; G33-F37; K50-F52; D58-R75; G91-A95; D101; N114; V118-E119; N125-H128; S139; V154-N156; R164-F167; F178-T185; K198-G200; N214; F219-P232; E239-S245; V250-N251; I256-P259; P266-E276; K290-I293 |
| CH65-EH - 5NFQ | Active site | D102, Y150, Y209, D249, H277 |
| | Buried amino acids | A9, P11, I14, L16-G22, A26, V28-R43, Q45-I46, F49-S50, F53-R64, S66, P69, Y75, P77, L80-I81, D83-F85, L87-A89, L91, I93, F96-L113, G115, R119-V120, R122-P131, I133-F134, L138-Y139, V143, E146-F154, A158-N159, L162-V163, G167, L171-L172, K174-V176, W178, M184, E188-R189, L192-L193, W196, A201-P213, M219, P222-F223, V225, T230, L234, L237, I239-A246, D249, A251-L252, P254, N256-L257, L260, I263-I264, L267, I269-V270, V272, C275-P284, V287-N288, A290-E292, F294-L295 |
| | Surface amino acids | M1-V8; L10; N12-G13; H15; P23-D25; P27; H44; R47-H48; D51-R52; G65; S67-K68; Q70-A74; T76; D78-K79; G82; L86; D89-T90; G92; G94-S95; G114; Q116-L118; E121; A132; Q135-L137; T140-P142; Q144-R145; R155-P157; D160-A161; K164-H166; L168-G170; M173; K177; D179-A183; E185-E187; D190-Q191; R194-D195; Q197-D200; I214-T218; D220-A221; K224; P226-Y229; P231-Q233; P235-R236; T238; L247-D248; L250; P253; E255; E258-G259; E261-E262; D265-P266; T268; R271; P273-D274; D285-A286; A289; F293; A296-H298 |
| | Main domain | M1-A132; P213-H298 |
| | Cap domain | I133-S212 |
| | Cap-loop | Q175-Q185 |
| | NC-loop | Q115-I133 |
| | Alpha-helices | H39-F49; P77-L91; W103-G115; F134-T140; P142-R155; P157-H166; L168-K174; P186-Q197; H199-A211; P254-I263; P284-L295 |
| | Beta-sheets | K6-L10; G13-E21; V28-H33; R54-P58; T97-H101; R122-A126; T241-A246; L267-V272 |
| | Loops | M1-L5; P11-N12; G22-P27; G34-S38; S50-F53; D59-T76; G92-F96; D102; Q116-E121; N127-I133; H141; D156; G167; E175-E185; N198; S212-P240; L247-P253; E258-G259; I264-P266; P273A283; A296-H298 |

**Supplementary Table S2**. Differences in median Schneider entropy values between the median entropy values of the selected proteins' compartments and remaining positions of the trimmed Multiple Sequence Alignment (MSA). Negative values indicate compartments with lower variability and positive values indicate compartments with higher variability in comparison to the remaining positions in the trimmed MSA. Differences in median distances values that are marked bold passed the statistical significance of the Epps-Singleton two-sample test (p-value <0.05).

| Compartment | msEH | hsEH | StEH1 | TrEH | bmEH | Sibe-EH | CH65-EH |
|---|---|---|---|---|---|---|---|
| Active site | **-0.462** | **-0.462** | **-0.462** | **-0.462** | **-0.462** | **-0.462** | **-0.462** |
| Buried amino acids | **-0.066** | **-0.091** | **-0.113** | **-0.026** | **-0.087** | **-0.095** | **-0.080** |
| Surface amino acids | **0.207** | **0.207** | **0.197** | **0.189** | **0.219** | **0.234** | **0.219** |
| Main domain | **0.074** | **0.079** | **0.015** | **0.080** | **0.049** | **0.018** | **0.043** |
| Cap domain | **0.111** | **0.095** | **0.128** | **0.108** | **0.131** | **0.164** | **0.144** |
| Cap-loop | **0.082** | **0.043** | **0.111** | **0.100** | **0.094** | **0.226** | **0.162** |
| NC-loop | **0.081** | **0.081** | **0.081** | **0.081** | **0.050** | **0.004** | **-0.007** |
| Helices | **0.117** | **0.109** | **0.106** | **0.126** | **0.112** | **0.109** | **0.091** |
| Loops | **0.052** | 0.055 | 0.054 | **0.037** | **0.055** | **0.097** | **0.110** |
| Strands | **0.007** | **-0.008** | **-0.014** | **-0.002** | **-0.011** | **-0.006** | **-0.015** |

**Supplementary Table S3**. List of corresponding tunnels identified in both the crystal structure and in the MD simulation for each analysed protein structure.

| CRank | MDRank | Name |
|---|---|---|
| msEH - 1CQZ (resolution 2.80 Å) | | |
| 1 | 1 | Tc/m1 |
| 2 | 2 | Tm1 |
| 3 | 6 | Tcap4 |
| 4 | 5 | Tm2 |
| 5 | 3 | Tm3 |
| 6 | 9 | Tside |
| hsEH - 1S8O (resolution 2.60 Å) | | |
| 1 | 2 | Tc/m1 |
| 2 | 1 | Tm1 |
| 3 | 8 | Tg |
| 4 | 15 | Tc/m2 |
| 5 | 5 | Tm5 |
| 6 | 16 | Tcap4 |
| 7 | 9 | Tcap2 |
| 8(9)* | 14 | Tm3 |
| StEH1 - 2CJP (resolution 1.95 Å) | | |
| 1 | 1 | Tm1 |
| 2 | 3 | Tc/m1 |
| 3 | 2 | Tm2 |
| 4 | 12 | Tcap3 |
| 5 | 7 | Tcap6 |
| 6 | 16 | Tc/m_back |
| 7 | 9 | Tcap5 |
| TrEH - 5URO (resolution 1.70 Å) | | |
| 1 | 1 | Tc/m1 |
| 2 | 2 | Tm1 |

| | | |
|---|---|---|
| 3 | 3 | Tcap4 |
| 4 | 4 | Tside |
| 5 | 6 | Tm5 |
| 7 | 13 | Tback |
| bmEH - 4NZZ (resolution 1.75 Å) | | |
| 2(1)** | 1 | Tc/m1 |
| 3 | 3 | Tc/m_back |
| 4 | 4 | Tcap7 |
| Sibe-EH - 5NG7 (resolution 1.37 Å) | | |
| 1 | 1 | Tc/m1 |
| 2 | 3 | Tc/m3 |
| 3 | 9 | Tc/m_back |
| 4 | 6 | Tcap4 |
| 5 | 12 | Tc/m2 |
| 6 | 13 | Tside |
| CH65-EH - 5NFQ (resolution 1.60 Å) | | |
| 1 | 2 | Tc/m2 |
| 2 | 1 | Tc/m_back |
| 3 | 5 | Tc/m_side |

(*) - two tunnels identified in the crystal structure of hsEH (8th and 9th) correspond to the 14th tunnel identified during the MD simulation. We chose the 8th tunnel from the crystal structure due to the higher similarity rate.
(**) - two tunnels identified in the crystal structure of bmEH (1st and 2nd) corresponded to the 1st tunnel identified during the MD simulation. We chose the 2nd tunnel from the crystal structure due to the higher similarity rate.

**Supplementary Figure S1**. Issue related to identification of asymmetrical tunnels based on the example of the Tc/m tunnels identified by CAVER software during MD simulations.

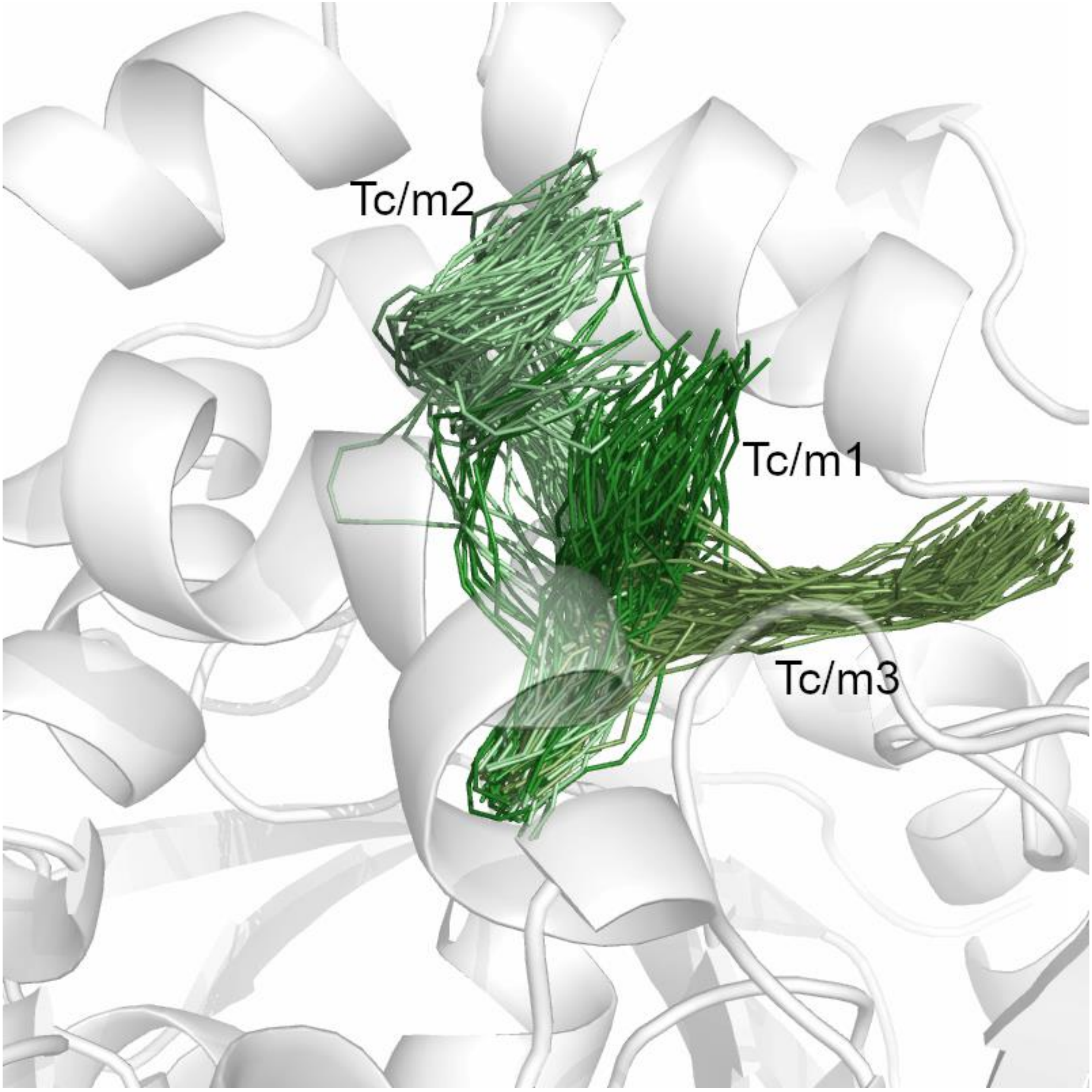

**Supplementary Table S4.** Comparison of maximal bottleneck radii measured in corresponding tunnels identified in both the crystal structure (CR) and in the MD simulation (MD) for each protein structure.

| CRank | MDRank | Name | CR max bottleneck | MD max bottleneck |
|---|---|---|---|---|
| msEH - 1CQZ (resolution 2.80 Å) | | | | |
| 1 | 1 | Tc/m1 | 2.1 | 3.2 |
| 2 | 2 | Tm1 | 2.0 | 3.1 |
| 3 | 6 | Tcap4 | 1.1 | 1.9 |
| 4 | 5 | Tm2 | 1.0 | 2.2 |
| 5 | 3 | Tm3 | 1.0 | 2.4 |
| 6 | 9 | Tside | 1.0 | 1.2 |
| hsEH - 1S8O (resolution 2.60 Å) | | | | |
| 1 | 2 | Tc/m1 | 1.6 | 2.7 |
| 2 | 1 | Tm1 | 1.8 | 2.9 |
| 3 | 8 | Tg | 1.1 | 1.9 |
| 4 | 15 | Tc/m2 | 1.3 | 2.0 |
| 5 | 5 | Tm5 | 1.2 | 1.8 |
| 6 | 16 | Tcap4 | 1.1 | 1.3 |
| 7 | 9 | Tcap2 | 1.0 | 1.7 |
| 8 | 14 | Tm3 | 0.9 | 1.1 |
| StEH1 - 2CJP (resolution 1.95 Å) | | | | |
| 1 | 1 | Tm1 | 1.8 | 3.1 |
| 2 | 3 | Tc/m1 | 1.4 | 2.1 |
| 3 | 2 | Tm2 | 1.8 | 2.7 |
| 4 | 12 | Tcap3 | 1.1 | 1.4 |
| 5 | 7 | Tcap6 | 1.0 | 1.4 |
| 6 | 16 | Tc/m_back | 1.1 | 1.3 |
| 7 | 9 | Tcap5 | 0.9 | 1.4 |
| TrEH - 5URO (resolution 1.70 Å) | | | | |
| 1 | 1 | Tc/m1 | 2.4 | 2.9 |
| 2 | 2 | Tm1 | 1.5 | 2.7 |
| 3 | 3 | Tcap4 | 1.3 | 2.0 |

| | | | | |
|---|---|---|---|---|
| 4 | 4 | Tside | 1.0 | 2.3 |
| 5 | 6 | Tm5 | 0.9 | 1.8 |
| 7 | 13 | Tback | 0.9 | 1.1 |
| bmEH - 4NZZ (resolution 1.75 Å) | | | | |
| 2 | 1 | Tc/m1 | 1.9 | 2.7 |
| 3 | 3 | Tc/m_back | 1.1 | 1.9 |
| 4 | 4 | Tcap7 | 0.9 | 1.2 |
| Sibe-EH - 5NG7 (resolution 1.37 Å) | | | | |
| 1 | 1 | Tc/m1 | 1.9 | 2.5 |
| 2 | 3 | Tc/m3 | 1.1 | 1.9 |
| 3 | 9 | Tc/m_back | 1.2 | 1.2 |
| 4 | 6 | Tcap4 | 1.1 | 1.7 |
| 5 | 12 | Tc/m2 | 0.9 | 1.4 |
| 6 | 13 | Tside | 0.9 | 1.2 |
| CH65-EH - 5NFQ (resolution 1.60 Å) | | | | |
| 1 | 2 | Tc/m2 | 1.4 | 2.3 |
| 2 | 1 | Tc/m_back | 1.6 | 2.5 |
| 3 | 5 | Tc/m_side | 1.2 | 1.5 |

**Supplementary Figure S2.** Correlation between maximal bottleneck radii measured in corresponding tunnels identified in both the crystal structure and in the MD simulation for each protein structure.

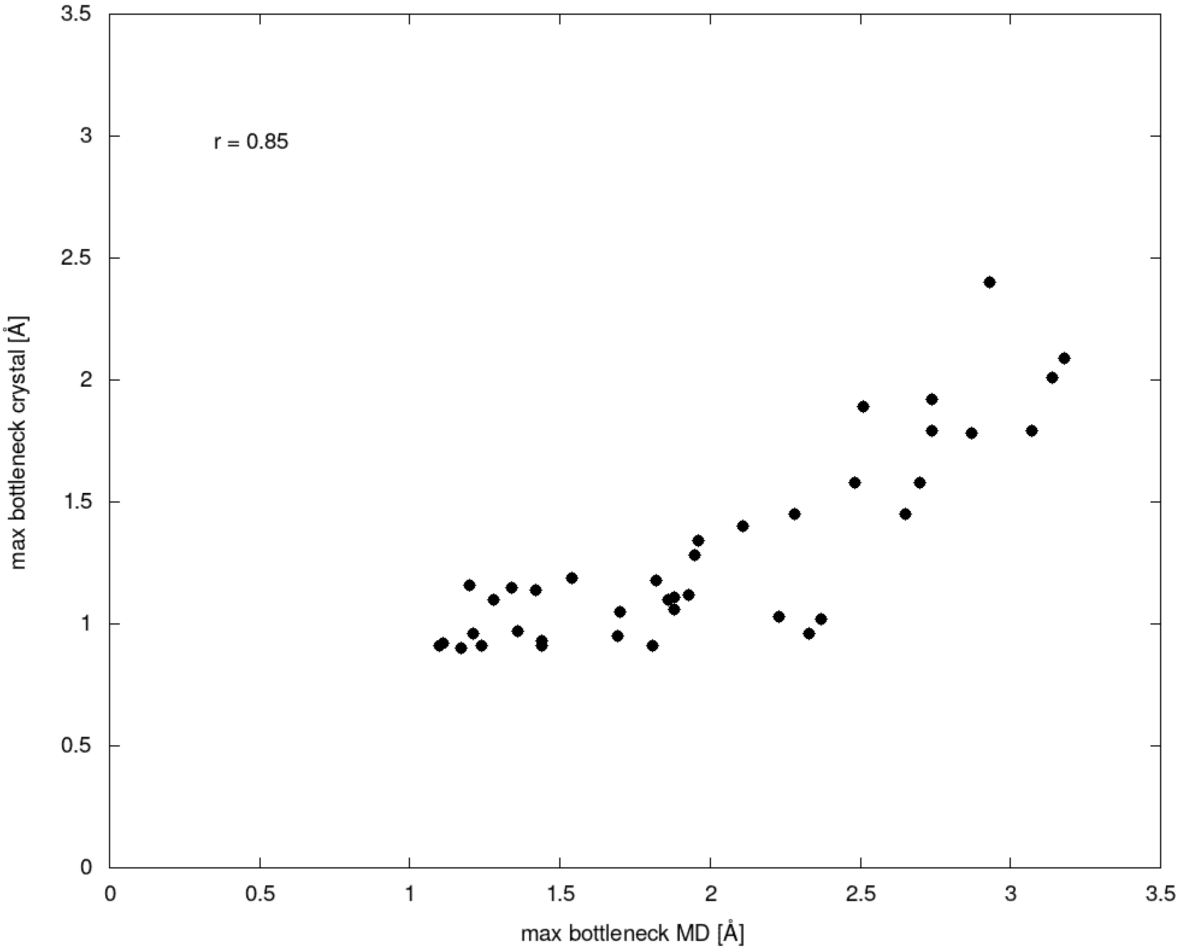

**Supplementary Table S5.** List of amino acids forming analysed tunnels in msEH structure.

| MDRank | Name | Tunnel-lining residues | Tunnel-lining residues without the surface residues | Surface tunnel-lining residues |
|---|---|---|---|---|
| 1 | Tc/m1 | F265, P266, D333, Y381, F385, S405, F406, R408, F415, I416, V418, Y465, D495, I496, V497, H524, W524 | F265, P266, D333, Y381, F385, F406, V418, Y465, D495, V497, H524, W524 | S405, R408, F415, I416, I496 |
| 2 | Tm1 | F265, P266, H332, D333, W334, V337, N357, T358, P359, F360, M361, P362, P363, D364, V367, S368, P369, M370, V372, I373, I376, F379, Y381, Q382, F385, F406, I416, V418, Y465, T468, W472, V497, L498, M502, H523, W524 | F265, P266, H332, D333, W334, V337, N357, T358, P359, F360, M361, P362, D364, V367, I373, I376, F379, Y381, Q382, F385, F406, V418, Y465, T468, W472, V497, L498, M502, H523, W524 | P363, S368, P369, M370, V372, I416 |
| 5 | Tm2 | F265, P266, M308, H322, D333, W334, V337, W340, N341, L344, F345, T358, P359, F360, M361, P362, P363, D364, P369, V372, I373, F379, Y381, Q382, F385, F406, I416, V418, Y465, T468, N471, W472, W474, S475, C476, K477, G478, L479, G480, R481, K482, I483, V497, L498, M502, K508, W509, H523, W524 | F265, P266, M308, H322, D333, W334, V337, W340, N341, L344, T358, P359, F360, M361, P362, D364, I373, F379, Y381, Q382, F385, F406, V418, Y465, T468, W472, C476, G480, I483, V497, L498, M502, W509, H523, W524 | F345, P363, P369, V372, I416, N471, W474, S475, K477, G478, L479, R481, K482, K508 |
| 3 | Tm3 | G264, F265, P266, E267, H332, D333, W334, C337, L356, N357, T358, P359, F379, Y381, Q382, F385, F406, F415, I416, V418, I427, Y465, M489, V490, T491, A492, E493, K494, D495, I496, V497, L498, R499, P500, E501, M502, S503, K504, R515, H517, C521, G522, H523, W524 | G264, F265, P266, E267, H332, D333, W334, C337, L356, N357, T358, P359, F379, Y381, Q382, F385, F406, V418, I427, Y465, M489, V490, T491, A492, D495, V497, L498, P500, M502, S503, R515, C521, G522, H523, W524 | F415, I416, E493, K494, I496, R499, E501, K504, H517 |
| 9 | Tside | G264, F265, P266, E267, S268, F270, S271, W272, R273, Y274, H332, D333, W334, V337, L356, N357, T358, P359, Y381, Q382, F385, F406, F407, R408, I416, V418, K439, I440, T441, T442, E445, I446, Y449, Y465, D495, I496, V497, L498, G522, H523, W524, T525, Q526, I527, E528 | G264, F265, P266, E267, S268, F270, S271, W272, H332, D333, W334, V337, L356, N357, T358, P359, Y381, Q382, F385, F406, F407, V418, I440, T441, E445, I446, Y449, Y465, D495, V497, L498, G522, H523, W524, T525, Q526, I527, E528 | R273, Y274, R408, I416, K439, T442, I496 |
| 6 | Tcap4 | G264, F265, P266, M308, H332, D333, W334, V337, N357, T358, P359, F360, M361, P362, P363, S368, P369, M370, V372, I373, R374, F379, Y381, Q382, L383, F385, Q386, F406, I416, V418, W464, Y465, R466, N467, T468, E469, R470, N471, W472, D495, I496, V497, L498, M502, H523, W524 | G264, F265, P266, M308, H332, D333, W334, V337, N357, T358, P359, F360, M361, P362, I373, F379, Y381, Q382, L383, F385, F406, V418, W464, Y465, N467, T468, W472, D495, V497, L498, M502, H523, W524 | P363, S368, P369, M370, V372, R374, Q386, I416, R466, E469, R470, N471, I496 |

**Supplementary Table S6**. List of amino acids forming analysed tunnels in hsEH structure.

| MDRank | Name | Tunnel-lining residues | Tunnel-lining residues without the surface residues | Surface tunnel-lining residues |
|---|---|---|---|---|
| 2 | Tc/m1 | F267, P268, D335, W336, Y383, Q384, F387, S407, L408, F409, R410, A411, S412, S415, V416, L417, S418, M419, Y466, K495, D496, F497, V498, L499, G523, H524, W525 | F267, P268, D335, W336, Y383, Q384, F387, L408, F409, V416, M419, Y466, D496, V498, L499, G523, H524, W525 | S407, R410, A411, S412, S415, L417, S418, K495, F497 |
| 15 | Tc/m2 | G266, F267, P268, H334, D335, W336, V380, F381, Y383, Q384, F387, S407, L408, F409, R410, A411, S412, D413, E414, S415, V416, L417, S418, M419, H420, K421, V422, A425, L428, V430, N431, S432, Y466, D496, F497, V498, L499, H524, W525 | G266, F267, P268, H334, D335, W336, F381, Y383, Q384, F387, L408, F409, V416, M419, V422, A425, L428, S432, Y466, D496, V498, L499, H524, W525 | V380, S407, R410, A411, S412, D413, E414, S415, L417, S418, H420, K421, V430, N431, F497 |
| 8 | Tg | G266, F267, P268, D335, W336, P379, V380, F381, D382, Y383, Q384, F387, L408, S415, L417, S418, M419, H420, Y466, D496, F497, V498, L499, H524, W525 | G266, F267, P268, D335, W336, F381, D382, Y383, Q384, F387, L408, M419, Y466, D496, V498, L499, H524, W525 | P379, V380, S415, L417, S418, H420, F497 |
| 1 | Tm1 | F267, P268, D335, W336, M339, T360, P361, I363, P371, S374, I375, F381, Y383, Q384, L408, M419, Y466, M469, F497, V498, L499, M503, H524, W525 | F267, P268, D335, W336, M339, T360, P361, I363, I375, F381, Y383, Q384, L408, M419, Y466, M469, V498, L499, M503, H524, W525 | P371, S374, F497 |
| 14 | Tm3 | F267, P268, D335, W336, M339, N359, T360, P361, I363, P381, Y383, Q384, L408, M419, Y466, M469, M490, V491, T492, A493, E494, K495, D496, P497, V498, L499, V500, P501, Q502, M503, S504,H524, W525 | F267, P268, D335, W336, M339, N359, T360, P361, I363, P381, Y383, Q384, L408, M419, Y466, M469, M490, V491, T492, A493, D496, V498, L499, V500, P501, M503, S504,H524, W525 | E494, K495, F497, Q502 |
| 9 | Tcap2 | F267, P268, D335, W336, Y383, Q384, Y386, F387, A393, E396, L397, E398, Q399, N400, R403, T404, L408, L417, M419, G427, L428, P429, V430, S432, F459, L463, Y466, D496, F497, V498, L499, H524, W525 | F267, P268, D335, W336, Y383, Q384, Y386, F387, A393, E396, L397, T404, L408, M419, L428, P429, S432, F459, L463, Y466, D496, V498, L499, H524, W525 | E398, Q399, N400, R403, L417, G427, V430, F497 |
| 16 | Tcap4 | D335, Q384, M339, L372, I375, Y383, Q384, Q388, Y466, M469, V498, L499, H524 | D335, Q384, M339, L372, I375, Y383, Q384, Q388, Y466, M469, V498, L499, H524 | - |

**Supplementary Table S7**. List of amino acids forming analysed tunnels in StEH1 structure.

| MDRank | Name | Tunnel-lining residues | Tunnel-lining residues without the surface residues | Surface tunnel-lining residues |
|---|---|---|---|---|
| 3 | Tc/m1 | F33, P34, D105, W106, L109, S129, V130, L145, Y149, H153, Y154, I155, I180, P186, A187, P188, F189, Y235, D265, L266, V267, Y268, H269, I270, P271, H300, F301 | F33, P34, D105, W106, L109, S129, V130, L145, Y149, H153, Y154, I155, I180, Y235, D265, V267, Y268, I270, H300, F301 | P186, A187, P188, F189, L266, H269, P271 |
| 16 | Tc/m_back | F33, P34, G61, Y62, L70, D105, Y154, F158, F168, V176, L177, I180, P189, F191, F223, H226, G227, F228, T229, G230, A231, V232, Y235, L266, V267, H300, H301 | F33, P34, G61, Y62, D105, Y154, F158, F168, V176, L177, I180, F191, F223, F228, G230, A231, V232, Y235, V267, H300, H301 | L70, P189, H226, G227, T229, L266 |
| 1 | Tm1 | F33, P34, D105, W106, L109, S129, V130, H131, F132, S133, K134, N136, V141, G144, L145, I148, H153, Y154, I155, F158, I180, P189, Y235, L238, D265, L266, V267, I270, P271, A273, H300, F301 | F33, P34, D105, W106, L109, S129, V130, H131, F132, S133, N136, V141, L145, H153, Y154, I155, F158, I180, Y235, L238, D265, V267, I270, A273, H300, F301 | K134, G144, I148, P189, L266, P271 |
| 12 | Tcap3 | F33, P34, D105, Y154, F158, I164, E165, F168, A169, V176, I180, F189, F191, G227, F228, V232, Y235, L266, V267, H300, F301 | F33, P34, D105, Y154, F158, I164, F168, V176, I180, F191, F228, V232, Y235, V267, H300, F301 | E165, A169, F189, G227, L266 |
| 9 | Tcap5 | F33, P34, D105, W106, Y154, I155, F158, F168, S175, V176, K178, L179, I180, L181, T182, Y183, R184, F189, F191, L197, I200, P201, D202, A203, V205, Y235, L266, V267, H300, F301 | F33, P34, D105, W106, Y154, I155, F158, F168, V176, L179, I180, L181, F191, L197, I200, P201, Y235, V267, H300, F301 | S175, K178, T182, Y183, R184, F189, D202, A203, V205, L266 |
| 7 | Tcap6 | F33, P34, D105, W106, Y154, K178, I180, L181, T182, Y183, R184, D185, F189, A203, V205, L207, S208, S209, W210, L211, S212, E213, L216, Y235, D265, L266, V267, A299, H300, F301, E305 | F33, P34, D105, W106, Y154, I180, L181, W210, L211, L216, Y235, D265, V267, A299, H300, F301, E305 | K178, T182, Y183, R184, D185, F189, A203, V205, L207, S208, S209, S212, E213, L266 |

**Supplementary Table S8.** List of amino acids forming analysed tunnels in TrEH structure.

| MDRank | Name | Tunnel-lining residues | Tunnel-lining residues without the surface residues | Surface tunnel-lining residues |
|---|---|---|---|---|
| 1 | Tc/m1 | W46, P47, D116, Y167, M193, F205, T207, D286, N287, A288, L289, H313, W314 | W46, P47, D116, Y167, M193, F205, D286, A288, L289, H313, W314 | T207, N287 |
| 2 | Tm1 | T46, P47, D116, W117, A120, T141, H144, F165, Y167, Q168, M193, F205, T207, Y252, A288, L289, H313, W314 | T46, P47, D116, W117, A120, T141, H144, F165, Y167, Q168, M193, F205, Y252, A288, L289, H313, W314 | T207 |
| 6 | Tm5 | G45, T46, P47, T88, L89, K90, S91, H115, D116, W117, G118, G119, A120, V121, V122, W123, R124, T125, A126, Y127, Y128, H129, P130, V136, 138, C140, T141, P142, L143, H144, P145, L146, S147, L153, M162, N164, F165, Y167, Q168, L171, M193, F205, T207, W251, Y252, R255, L257, N258, A259, K260, D261, E262, M263, D264, R265, A266, P270, P271, L272, R273, F274, E275, M276, D286, N287, A288, L289, M293, F300, Y301, H313, W314 | G45, T46, P47, T88, L89, K90, S91, H115, D116, W117, G118, G119, A120, V121, V122, W123, R124, T125, A126, Y127, H129, V136, 138, C140, T141, P142, L143, H144, P145, L153, M162, N164, F165, Y167, Q168, L171, M193, F205, W251, Y252, R255, A259, M263, P271, F274, M276, D286, A288, L289, M293, F300, Y301, H313, W314 | Y128, P130, L146, S147, T207, L257, N258, K260, D261, E262, D264, R265, A266, P270, L272, R273, E275, N287 |
| 4 | Tside | G45, T46, P47, D48, G52, R54, H115, D116, Y167, M193, F194, F205, L224, L225, Y233, A288, H313, W314, L316, T317 | G45, T46, P47, D48, G52, R54, H115, D116, Y167, M193, F194, F205, L224, L225, Y233, A288, H313, W314, L316, T317 | |
| 13 | Tback | E16, T17, K18, Q19, I20, K23, T24, T25, S26, Y27, L29, V41, L42, V43, H44, G45, W46, P47, D48, W53, V68, A69, P70, N71, M72, L73, Y75, T78, N86, F87, T88, L89, S91, V92, S93, A94, D95, I96, A97, E98, L99, A100, S102, F103, H115, D116, W117, G118, G119, A120, V121, R124, C140, T141, P142, L143, H144, P145, S147, Y150, L153, M162, N164, F165, Y167, Q168, L171, M193, F205, T207, G247, P248, N250, W251, Y252, R253, T254, R255, E256, N258, A259, E262, D286, N287, A288, L289, M293, H313, T314 | E16, T17, K18, Q19, I20, K23, T24, T25, S26, Y27, L29, V41, L42, V43, H44, G45, W46, P47, D48, W53, V68, A69, P70, N71, M72, L73, Y75, T78, N86, F87, T88, L89, S91, V92, S93, A94, D95, I96, A97, E98, L99, A100, F103, H115, D116, W117, G118, G119, A120, V121, R124, C140, T141, P142, L143, H144, P145, Y150, L153, M162, N164, F165, Y167, Q168, L171, M193, F205, G247, P248, N250, W251, Y252, R253, T254, R255, E256, A259, D286, A288, L289, M293, H313, T314 | S102, S147, T207, N258, E262, N287 |
| 3 | Tcap4 | T46, P47, L89, D116, W117, G118, G119, A120, T141, P142, H144, Y150, P152, L153, E154, M162, N164, F165, Y167, Q168, L169, L171, K172, M193, F205, T207, W251, Y252, R253, T254, R255, E256, N258, D286, A288, L289, M293, H313, W314 | T46, P47, L89, D116, W117, G118, G119, A120, T141, P142, H144, Y150, L153, M162, N164, F165, Y167, Q168, L169, L171, M193, F205, W251, Y252, R253, T254, R255, E256, D286, A288, L289, M293, H313, W314 | P152, E154, K172, T207, N258 |

**Supplementary Table S9**. List of amino acids forming analysed tunnels in bmEH structure.

| MDRank | Name | Tunnel-lining residues | Tunnel-lining residues without the surface residues | Surface tunnel-lining residues |
|---|---|---|---|---|
| 1 | Tc/m1 | F30, P31, D32, D97, W98, A141, Y144, F148, L168, V169, Y203, D239, P240, T241, F242, H267 | F30, P31, D32, D97, W98, A141, Y144, F148, L168, V169, Y203, D239, T241, F242, H267 | P240 |
| 3 | Tc/m_back | F30, P31, E71, I72, D97, W98, G101, G122, P123, F128, Y144, m145, L168, Y203, N205, L206, K207, R216, L219, F220, P221, T241, F242, H267 | F30, P31, E71, I72, D97, W98, G101, G122, P123, F128, Y144, m145, L168, Y203, N205, L206, K207, F220, T241, F242, H267 | R216, L219, P221 |
| 4 | Tcap7 | H28, G29, F30, P31, D32, F33, S34, I35, I36, W37, R38, G57, Y58, N59, K63, H96, D97, W98, Y144, M145, F148, Q154, M157, F158, F162, L165, L168, V169, I170, P172, G173, Y178, L179, T180, D182, D183, B184, Q185, A186, Y187, M188, N189, S190, W191, E192, N193, G194, S195, V196, L197, S198, M199, Y203, D239, P240, T241, F242, S266, H267, A268, Q270, H271, E272 | H28, G29, F30, P31, D32, F33, S34, I35, I36, W37, R38, G57, Y58, N59, K63, H96, D97, W98, Y144, M145, F148, Q154, M157, F162, L165, L168, V169, P172, G173, L179,182, D183, B184, A186, Y187, M188, W191, S195, V196, S198, M199, Y203, D239, T241, F242, S266, H267, A268, Q270, H271, E272 | E158, I170, Y178, T180, D182, Q185, N189, S190, E192, N193, G194, L197, P240 |

**Supplementary Table S10.** List of amino acids forming analysed tunnels in Sibe-EH structure.

| MDRank | Name | Tunnel-lining residues | Tunnel-lining residues without the surface residues | Surface tunnel-lining residues |
|---|---|---|---|---|
| 1 | Tc/m1 | F34, P35, D101, W102, K145, W147, Y148, F152, M173, L174, Q176, S177, Y209, D241, V242, A243, I244, H270, W271 | F34, P35, D101, W102, W147, Y148, F152, M173, L174, S177, Y209, D241, A243, I244, H270, W271 | K145, Q176, V242 |
| 12 | Tc/m2 | F34, P35, D101, W147, Y148, F152, L162, F169, L170, N172, M173, L174, S177, Y209, V242, A243, I244, H270, W271 | F34, P35, D101, W147, Y148, F152, L162, F169, L170, M173, L174, S177, Y209, A243, I244, H270, W271 | N172, V242 |
| 3 | Tc/m3 | F34, P35, H100, D101, W102, Y148, F152, M173, L174, Q176, S177, F178, V179, Y209, K240, D241, V242, A243, I244, C268, F269, H270, W271, E275 | F34, P35, H100, D101, W102, Y148, F152, M173, L174, S177, F178, Y209, D241, A243, I244, C268, F269, H270, W271, E275 | Q176, V179, K240, V242 |
| 9 | Tc/m_back | G33, F34, P35, L76, D101, W102, I105, N125, A126, P127, H128, P129, K130, A131, Y132, M133, T134, R135, T136, K137, N138, Q142, L143, K145, S146, Y148, V149, F150, F152, M173, S177, Y208, Y209, N212, L213, N214, D216, I217, I218, F219, S220, E221, K222, T223, V224, D241, V242, A243, I244, L248, H270, W271 | G33, F34, P35, L76, D101, W102, I105, N125, A126, P127, H128, P129, T136, L143, S146, Y148, V149, F150, F152, M173, S177, Y208, Y209, N212, L213, D216, I217, F219, K222, D241, A243, I244, L248, H270, W271 | K130, A131, Y132, M133, T134, R135, K137, N138, Q142, K145, N214, D216, I217, I218, S220, E221, T223, V224, V242 |
| 13 | Tside | G33, F34, P35, D36, Y39, V40, R42, F43, H100, D101, W102, Y148, F152, L170, K171, N172, M173, L174, I175, S177, F178, D182, L183, L184, T185, D188, L189, Y192, W196, Y209, D241, V242, A243, I244, H270, W271, Q273, L274, E275, P277 | G33, F34, P35, D36, Y39, V40, R42, F43, H100, D101, W102, Y148, F152, L170, M173, L174, S177, F178, L184, D188, L189, Y192, W196, Y209, D241, A243, I244, H270, W271, Q273, E275, P277 | K171, N172, I175, D182, L183, T185, V242, L274 |
| 6 | Tcap4 | F34, P35, D101, W102, I105, N125, A126, P127, Y132, T136, K137, Q142, L143, K145, S146, W147, Y148, V149, F150, Q153, M173, S177, Y209, L213, N214, P215, I218, F219, D241, V242, A243, I244, L248, H270, W271 | F34, P35, D101, W102, I105, N125, A126, P127, T136, L143, S146, W147, Y148, V149, F150, M173, S177, Y209, L213, F219, D241, A243, I244, L248, H270, W271 | Y132, K137, Q142, K145, Q153, N214, P215, I218, V242 |

**Supplementary Table S11**. List of amino acids forming analysed tunnels in CH65-EH structure.

| MDRank | Name | Tunnel-lining residues | Tunnel-lining residues without the surface residues | Surface tunnel-lining residues |
|---|---|---|---|---|
| 2 | Tc/m2 | D102, A147, S148, Y150, K177, W178, D179, Y209, D249, L250, A251, L252, H277, F278 | D102, A147, S148, Y150, W178, Y209, D249, A251, L252, H277, F278 | K177, D179, L250 |
| 5 | Tc/m_side | F35, D102, Y150, K177, W178, D179, R180, P181, S182, A183, M184, Y209, D248, D249, L250, A251, L252, C275, G276, H277, F278, W281, E282 | F35, D102, Y150, W178, M184, Y209, D249, A251, L252, C275, G276, H277, F278, W281, E282 | K177, D179, R180, P181, S182, A183, D248, L250 |
| 1 | Tc/m_back | F35, P77, D78, L80, I81, H101, D102, W103, G104, G105, A106, I107, W109, G110, L113, N127, A128, P129, H130, P131, A131, F134, L138, S148, Y150, I151, F154, W178, D179, Y208, Y209, S212, P213, I214, P232, L234, D249, A251, L252, H277, F278 | F35, P77, L80, I81, H101, D102, W103, G104, G105, A106, I107, W109, G110, L113, N127, A128, P129, H130, P131, F134, L138, S148, Y150, I151, F154, W178, Y208, Y209, S212, P213, L234, D249, A251, L252, H277, F278 | D78, A132, D179, I214, P232 |

**Supplementary Figure S3.** The distribution of the entropy values and the median entropy values of all tunnel-lining residues, tunnel-lining residues without the surface residues and the remaining positions of the trimmed MSA, and surface tunnel-lining residues (the violin and box plot) for the *M. musculus* soluble epoxide hydrolase (msEH). Statistically significant pairwise differences in the median distance values are marked by a star (*).

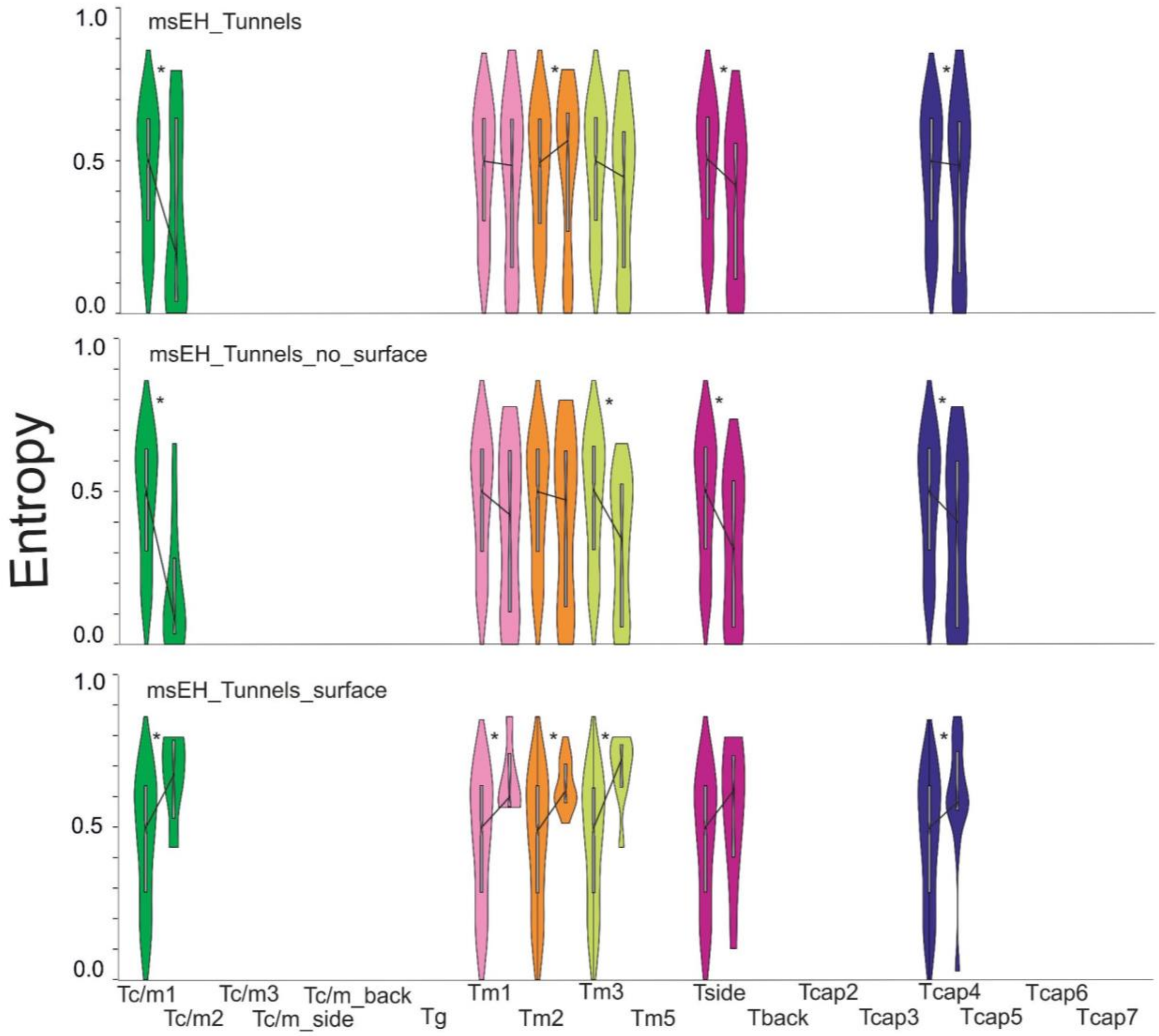

**Supplementary Figure S4**. The distribution of the entropy values and the median entropy values of all tunnel-lining residues, tunnel-lining residues without the surface residues and the remaining positions of the trimmed MSA, and surface tunnel-lining residues (the violin and box plot) for the *H. sapiens* soluble epoxide hydrolase (hsEH). Statistically significant pairwise differences in the median distance values are marked by a star (*).

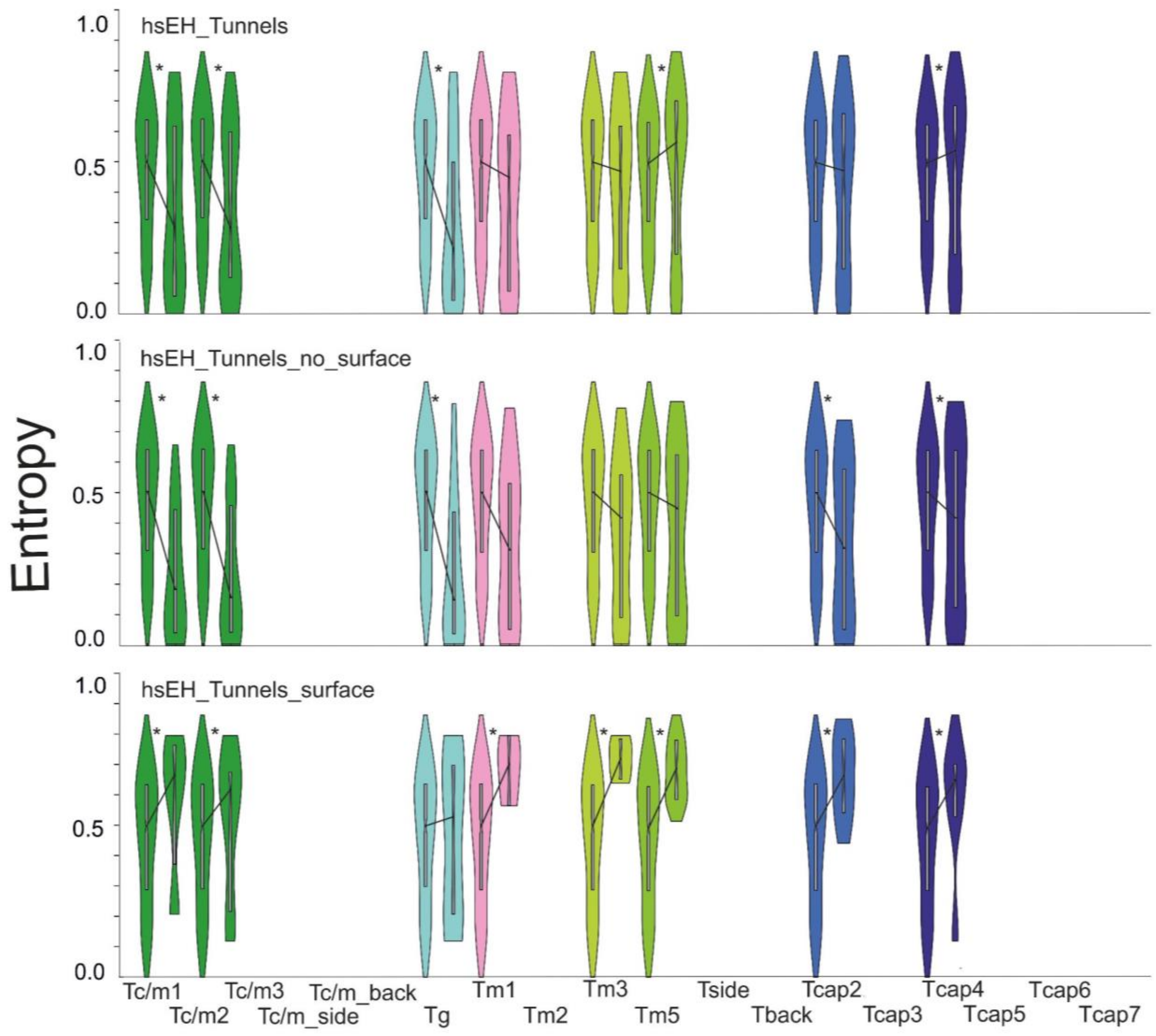

**Supplementary Figure S5.** The distribution of the entropy values and the median entropy values of all tunnel-lining residues, tunnel-lining residues without the surface residues and the remaining positions of the trimmed MSA, and surface tunnel-lining residues (the violin and box plot) for the *S. tuberosum* soluble epoxide hydrolase (StEH1). Statistically significant pairwise differences in the median distance values are marked by a star (*).

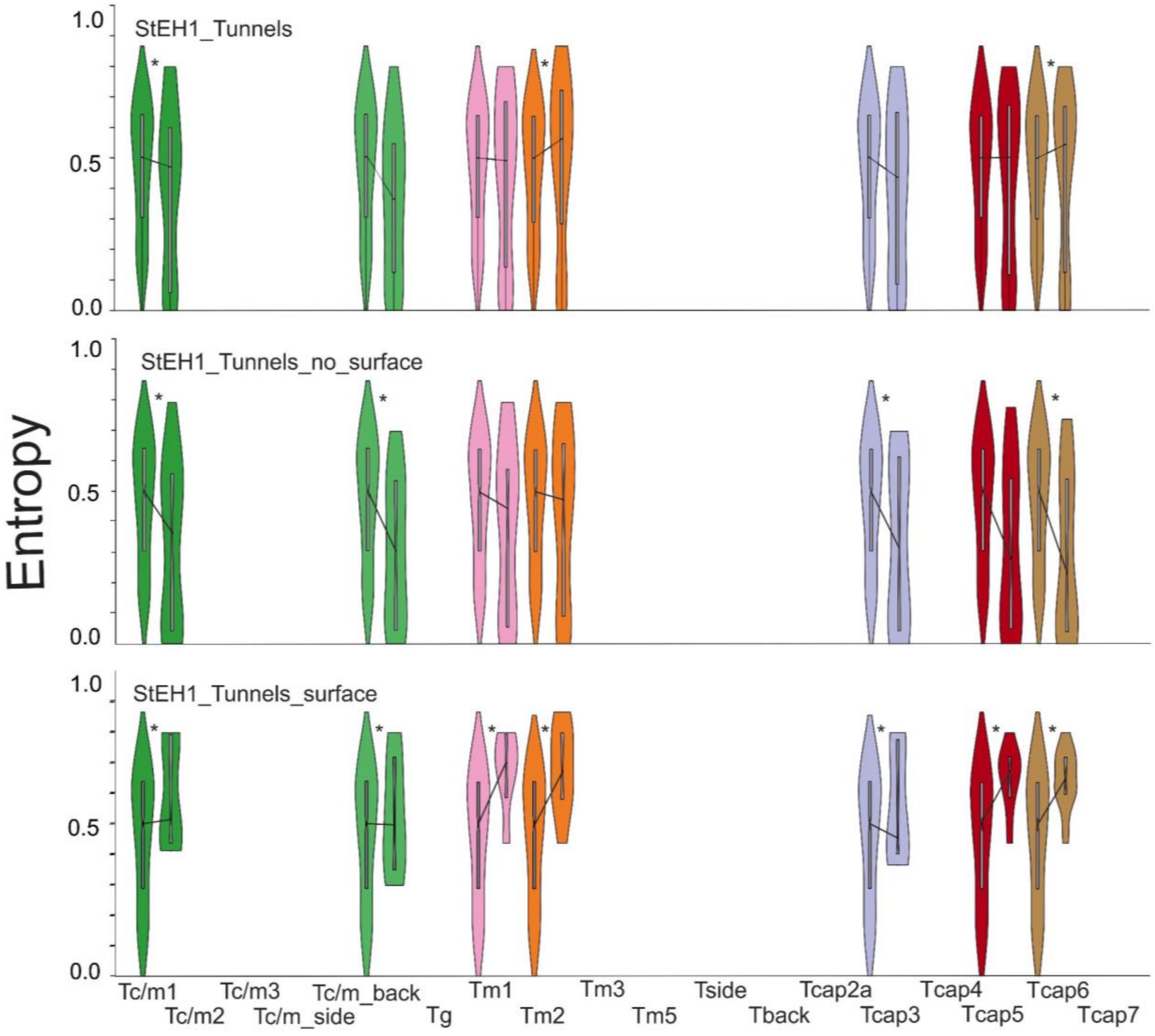

**Supplementary Figure S6.** The distribution of the entropy values and the median entropy values of all tunnel-lining residues, tunnel-lining residues without the surface residues and the remaining positions of the trimmed MSA, and surface tunnel-lining residues (the violin and box plot) for the *T. resei* soluble epoxide hydrolase (TrEH). Statistically significant pairwise differences in the median distance values are marked by a star (*). NA by the violin plots means that the number of surface residues was insufficient to obtain the p-value of the Epps–Singleton two-sample test. In the case of the Tside tunnel, no surface residues were identified.

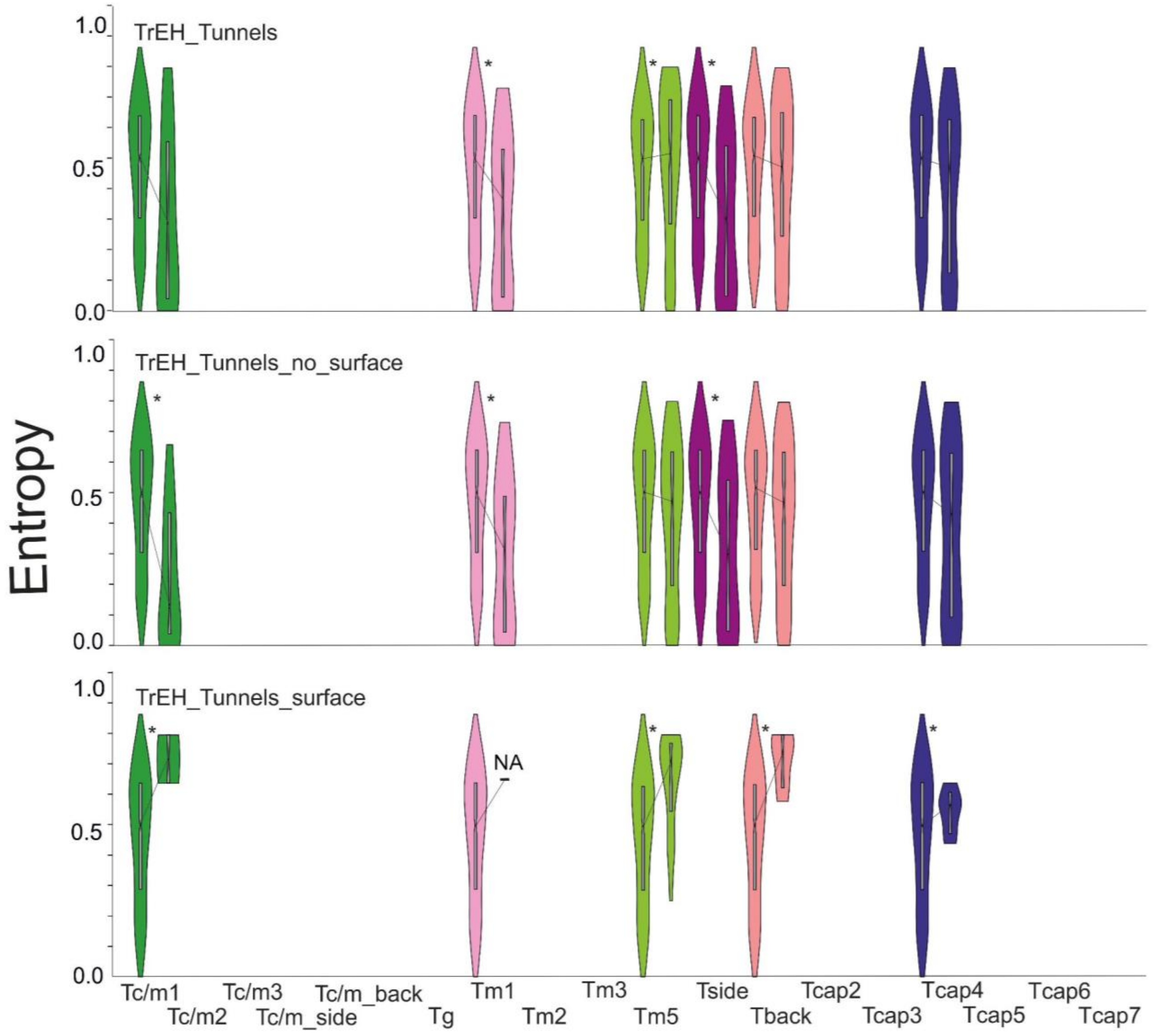

**Supplementary Figure S7.** The distribution of the entropy values and the median entropy values of all tunnel-lining residues, tunnel-lining residues without the surface residues and the remaining positions of the trimmed MSA, and surface tunnel-lining residues (the violin and box plot) for the *B. megaterium* soluble epoxide hydrolase (bmEH). Statistically significant pairwise differences in the median distance values are marked by a star (*). NA by the violin plots means that the number of surface residues was insufficient to obtain the p-value median distance of the Epps–Singleton two-sample test.

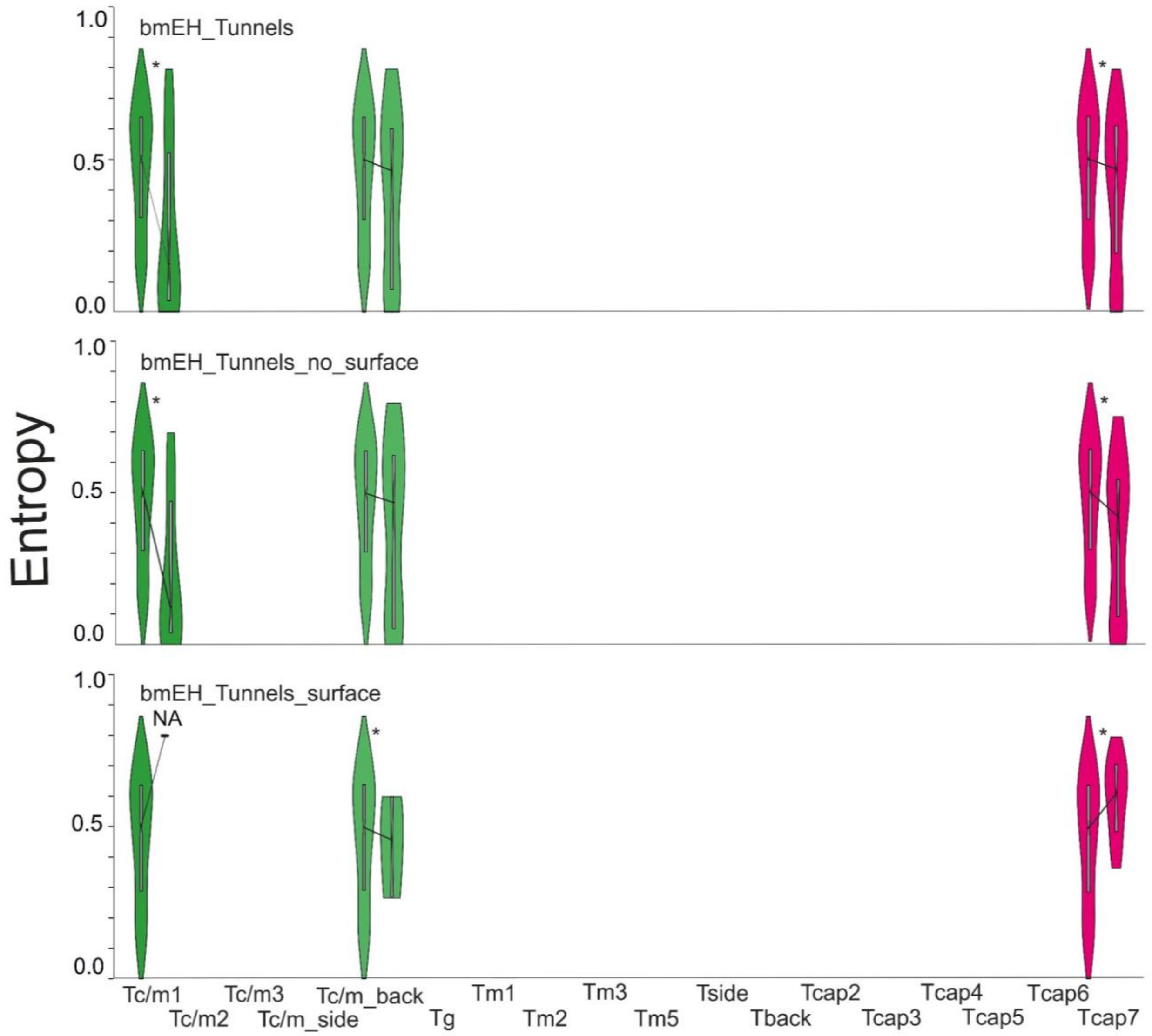

**Supplementary Figure S8.** The distribution of the entropy values and the median entropy values of all tunnel-lining residues, tunnel-lining residues without the surface residues and the remaining positions of the trimmed MSA, and surface tunnel-lining residues (the violin and box plot) for the thermophilic enzyme collected in hot springs in Russia (Sibe-EH). Statistically significant pairwise differences in the median distance values are marked by a star (*).

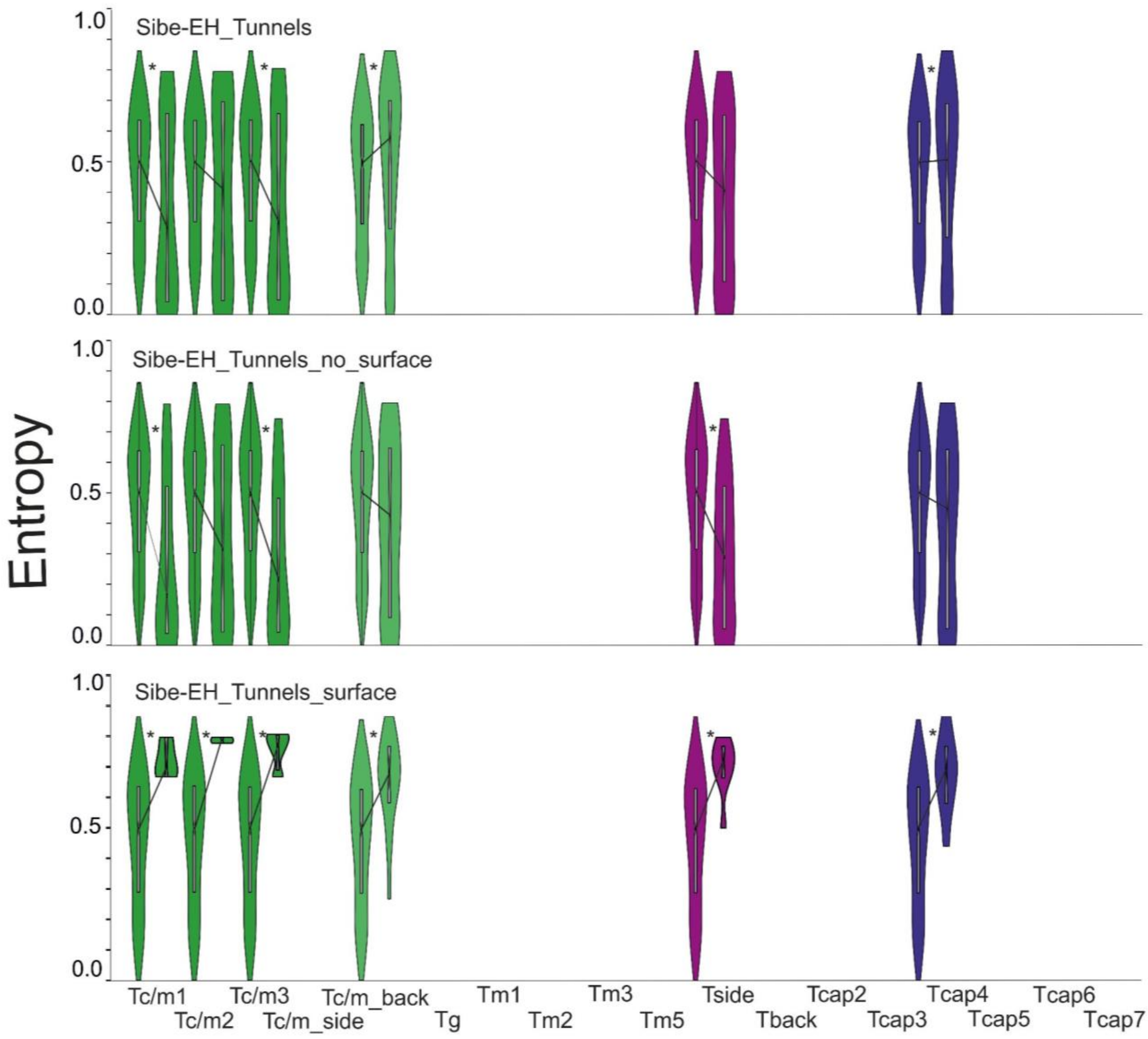

**Supplementary Figure S9.** The distribution of the entropy values and the median entropy values of all tunnel-lining residues, tunnel-lining residues without the surface residues and the remaining positions of the trimmed MSA, and surface tunnel-lining residues (the violin and box plot) for the thermophilic enzyme collected in hot springs in China (CH65-EH). Statistically significant pairwise differences in the median distance values are marked by a star (*).

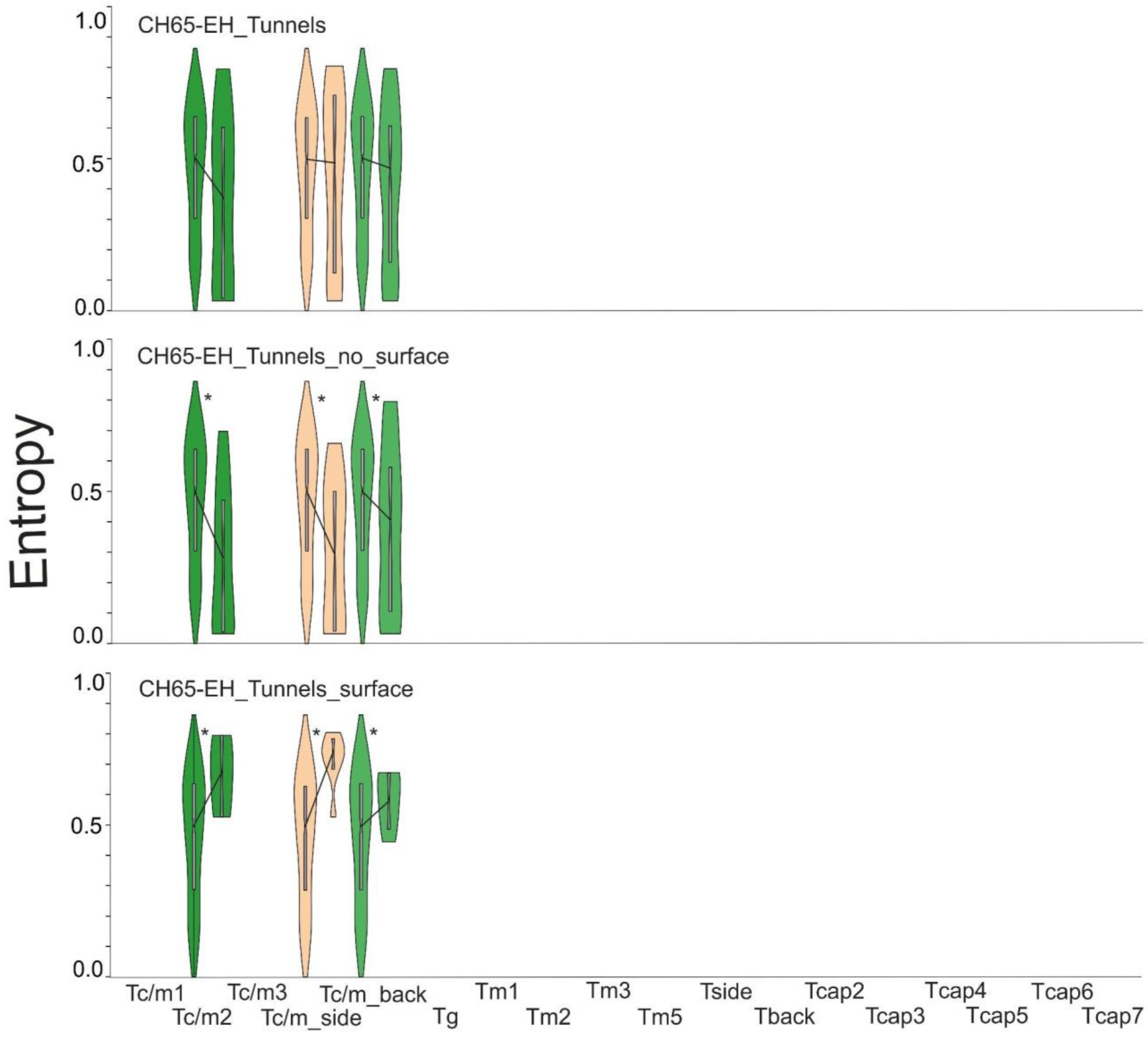

**Supplementary Table S12.** Differences in Schneider entropy values between the median distance of selected tunnel-lining residues and the median distances of the remaining positions of the trimmed Multiple Sequence Alignment (MSA). Negative values indicate compartments with lower variability and positive values indicate compartments with higher variability than the remaining positions of the trimmed MSA. Table consists of: the MD simulation tunnel ranking (MDRank), tunnel name, its average length, number of tunnel-lining residues, number of tunnel-lining residues without the surface residues, number of surface tunnel-lining residues, and the last three columns comprise of the differences between the median distances of particular tunnel-lining residues and the remaining positions of the trimmed MSA: median distances of tunnel-lining residues (7th column), median distances of the tunnel-lining residues without the surface residues (8th column), and median distances of the surface tunnel-lining residues (9th column). Differences between median distances values that are marked bold passed the statistical significance of the Epps-Singleton two-sample test (p-value <0.05). Hyphen (-) indicates that there were no surface residues. NA means that the number of surface residues was insufficient to obtain the p-value median distance of the Epps–Singleton two-sample test.

| MDRank | Name | Average length [Å] | Number of tunnel-lining residues | Number of tunnel-lining residues without the surface residues | Number of surface tunnel-lining residues | Tunnel-lining residues | Tunnel-lining residues without the surface residues | Surface tunnel-lining residues |
|---|---|---|---|---|---|---|---|---|
| msEH | | | | | | | | |
| 1 | Tc/m1 | 14 | 17 | 12 | 5 | **-0.307** | **-0.417** | **0.178** |
| 2 | Tm1 | 19 | 36 | 30 | 6 | -0.014 | -0.079 | **0.105** |
| 5 | Tm2 | 33 | 49 | 35 | 14 | **0.070** | -0.028 | **0.125** |
| 3 | Tm3 | 20 | 44 | 35 | 9 | -0.049 | **-0.165** | **0.226** |
| 9 | Tside | 32 | 45 | 38 | 7 | **-0.086** | **-0.192** | 0.125 |
| 6 | Tcap4 | 29 | 46 | 33 | 13 | **-0.014** | **-0.102** | **0.085** |
| hsEH | | | | | | | | |
| 2 | Tc/m1 | 13 | 27 | 18 | 9 | **-0.217** | **-0.321** | **0.169** |
| 15 | Tc/m2 | 21 | 39 | 24 | 15 | **-0.218** | **-0.349** | **0.121** |
| 8 | Tg | 13 | 25 | 18 | 7 | **-0.296** | **-0.357** | 0.029 |
| 1 | Tm1 | 14 | 24 | 21 | 3 | -0.053 | -0.189 | **0.204** |

| | | | | | | | | |
|---|---|---|---|---|---|---|---|---|
| 14 | Tm3 | 30 | 33 | 29 | 4 | -0.031 | -0.084 | **0.223** |
| 5 | Tm5 | 34 | 54 | 38 | 16 | **0.071** | -0.053 | **0.197** |
| 9 | Tcap2 | 23 | 33 | 25 | 8 | -0.027 | **-0.183** | **0.166** |
| 16 | Tcap4 | 25 | 13 | 12 | 1 | **0.040** | **-0.084** | **0.159** |
| StEH1 | | | | | | | | |
| 3 | Tc/m1 | 16 | 27 | 20 | 7 | **-0.031** | **-0.135** | **0.014** |
| 16 | Tc/m_back | 31 | 27 | 21 | 6 | -0.138 | **-0.197** | **-0.004** |
| 1 | Tm1 | 13 | 32 | 26 | 6 | -0.009 | -0.056 | **0.199** |
| 2 | Tm2 | 21 | 33 | 25 | 8 | **0.063** | -0.027 | **0.174** |
| 12 | Tcap3 | 24 | 21 | 16 | 5 | -0.065 | **-0.184** | **-0.046** |
| 9 | Tcap5 | 29 | 30 | 20 | 10 | 0.001 | -0.224 | **0.174** |
| 7 | Tcap6 | 28 | 31 | 17 | 14 | **0.045** | **-0.273** | **0.147** |
| TrEH | | | | | | | | |
| 1 | Tc/m1 | 5 | 13 | 11 | 2 | -0.216 | **-0.378** | **0.218** |
| 2 | Tm1 | 14 | 18 | 17 | 1 | **-0.132** | **-0.189** | 0.139(NA) |
| 6 | Tm5 | 32 | 72 | 54 | 18 | **0.017** | -0.030 | **0.216** |
| 4 | Tside | 24 | 20 | 20 | 0 | **-0.201** | **-0.201** | - |
| 13 | Tback | 52 | 89 | 83 | 6 | -0.037 | -0.049 | **0.243** |
| 3 | Tcap4 | 28 | 39 | 34 | 5 | -0.031 | -0.071 | **0.067** |
| bmEH | | | | | | | | |
| 1 | Tc/m1 | 6 | 16 | 15 | 1 | **-0.343** | **-0.378** | 0.297(NA) |

| | | | | | | | | |
|---|---|---|---|---|---|---|---|---|
| 3 | Tc/m_back | 27 | 24 | 21 | 3 | -0.037 | -0.031 | **-0.042** |
| 4 | Tcap7 | 35 | 63 | 50 | 13 | **-0.034** | **-0.082** | **0.114** |
| Sibe-EH | | | | | | | | |
| 1 | Tc/m1 | 10 | 19 | 16 | 3 | **-0.215** | **-0.343** | **0.200** |
| 12 | Tc/m2 | 18 | 19 | 17 | 2 | -0.090 | -0.187 | **0.288** |
| 3 | Tc/m3 | 13 | 24 | 20 | 4 | **-0.203** | **-0.292** | **0.277** |
| 9 | Tc/m_back | 30 | 52 | 33 | 19 | **0.082** | -0.076 | **0.188** |
| 13 | Tside | 27 | 40 | 32 | 8 | -0.097 | **-0.219** | **0.225** |
| 6 | Tcap4 | 22 | 35 | 26 | 9 | **0.009** | -0.053 | **0.194** |
| CH65-EH | | | | | | | | |
| 2 | Tc/m2 | 12 | 14 | 11 | 3 | -0.129 | **-0.216** | **0.178** |
| 5 | Tc/m_side | 24 | 23 | 15 | 8 | -0.011 | **-0.203** | **0.243** |
| 1 | Tc/m_back | 19 | 41 | 36 | 5 | -0.032 | **-0.093** | **0.088** |

**Supplementary Table S13.** Entropy values for selected tunnels, Tm1 from StEH1, Tc/m1 from hsEH, and Tc/m_back from bmEH. Surface amino acids are in italics. Active site residues are marked by an asterisks (*).

| StEH1 -Tm1 tunnel | | hsEH - Tc/m1 tunnel | | bmEH - Tc/m_back tunnel | |
|---|---|---|---|---|---|
| amino acid | Schneider entropy | amino acid | Schneider entropy | amino acid | Schneider entropy |
| F33 | 0.123 | F267 | 0.123 | F30 | 0.123 |
| P34 | <0.001 | P268 | <0.001 | P31 | <0.001 |
| D105* | 0.041 | D335* | 0.041 | E71 | 0.645 |
| W106 | 0.057 | W336 | 0.057 | I72 | 0.750 |
| L109 | 0.596 | Y383* | 0.037 | D97* | 0.041 |
| S129 | 0.417 | Q384 | 0.468 | W98 | 0.057 |
| V130 | 0.506 | F387 | 0.194 | G101 | 0.560 |
| H131 | 0.252 | L408 | 0.656 | G122 | 0.506 |
| F132 | 0.400 | *S407* | *0.775* | P123 | 0.252 |
| S133 | 0.729 | F409 | 0.539 | Y128 | 0.580 |
| *K134* | *0.632* | *R410* | *0.675* | T144 | 0.037 |
| N136 | 0.766 | *A411* | *0.666* | M145 | 0.468 |
| V141 | 0.564 | *S412* | *0.618* | L168 | 0.656 |
| *G144* | *0.701* | *S415* | *0.207* | Y203* | 0.032 |
| L145 | 0.554 | V416 | 0.209 | N205 | 0.601 |
| *I148* | *0.690* | *L417* | *0.527* | L206 | 0.776 |
| H153 | 0791 | *S418* | *0.215* | K207 | 0.780 |
| Y154* | 0.037 | M419 | 0.172 | *R216* | *0.598* |
| I155 | 0.468 | Y466* | 0.032 | *L219* | *0.266* |
| F158 | 0.194 | *K495* | 0.792 | F220 | 0.425 |
| I180 | 0.656 | D496* | 0.034 | *P221* | *0.455* |
| *F189* | *0.434* | *F497* | *0.795* | H241 | 0.473 |
| Y235* | 0.032 | V498 | 0.471 | F242 | 0.284 |
| L238 | 0.776 | L499 | 0.284 | H267* | 0.045 |
| D265* | 0.034 | G523 | 0.437 | | |
| *L266* | *0.795* | H524* | 0.045 | | |
| V267 | 0.472 | W525 | 0.312 | | |
| I270 | 0.560 | | | | |

| | |
|---|---|
| *P271* | *0.789* |
| A273 | 0.520 |
| H300* | 0.045 |
| F301 | 0.312 |

**Supplementary Figure S10.** The open (coloured green) and closed (coloured red) position of the F497 residue of hsEH. The protein is shown as cartoon and surface, and the F497 residue is shown as sticks.

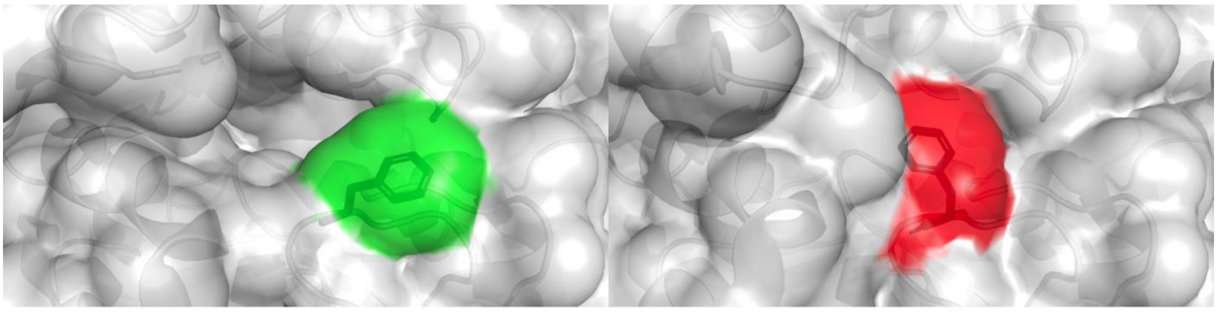

**Supplementary Table S13.** The list of parameters set for both CAVER plugin and CAVER 3.0 tunnels identification for each of the analysed systems.

| | shell_radius | shell_depth | probe_radius | clustering_threshold |
|---|---|---|---|---|
| 1CQZ | 4 | 6 | 0.9 | 5 |
| 1S8O | 4 | 6 | 0.9 | 4 |
| 2CJP | 4 | 5 | 0.9 | 4 |
| 5URO | 3 | 4 | 0.9 | 4 |
| 4NZZ | 4 | 6 | 0.9 | 5 |
| 5NG7 | 4 | 5 | 0.9 | 4 |
| 5NFQ | 5 | 5 | 0.9 | 4 |

**Supplementary Figure S10**. Representation of the created Multiple Sequence Alignment (MSA) of the epoxide hydrolases sequences. The red brace marks the sequences of the soluble epoxide hydrolases with known crystal structures. Gaps are marked in blue. MSA was pictured using the DECIPHER library for R (https://www.rdocumentation.org/packages/DECIPHER/versions/2.0.2).

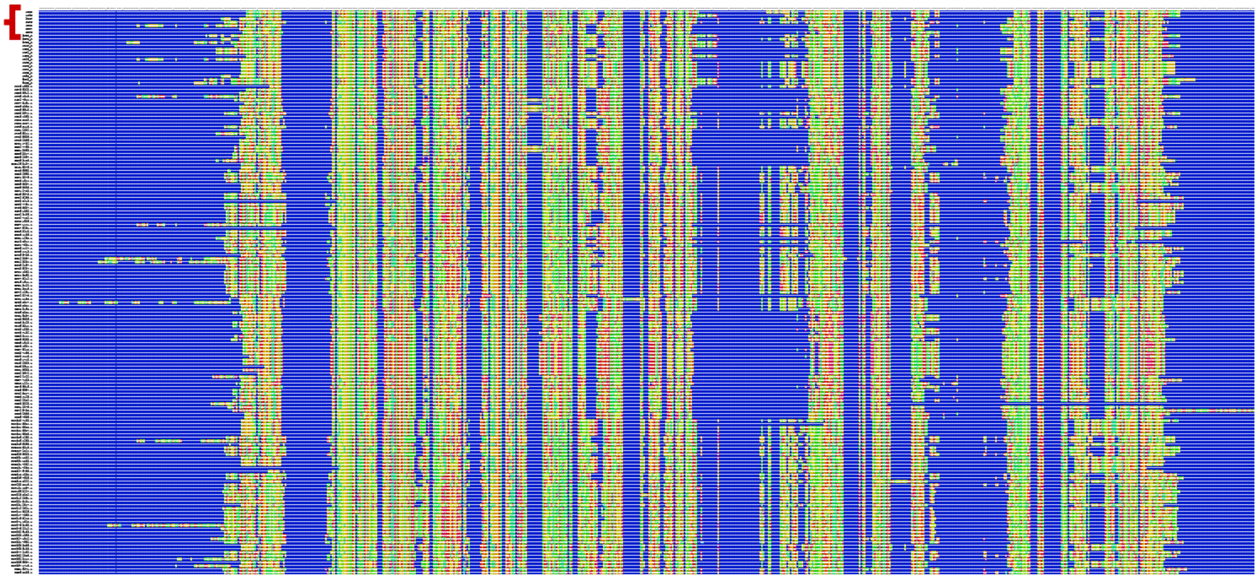

**Supplementary Figure S11**. Representation of the trimmed Multiple Sequence Alignment (MSA) of the epoxide hydrolases sequences. The red brace marks the sequences of the soluble epoxide hydrolases with known crystal structures. Gaps are marked in blue. MSA was pictured using the DECIPHER library for R (https://www.rdocumentation.org/packages/DECIPHER/versions/2.0.2).

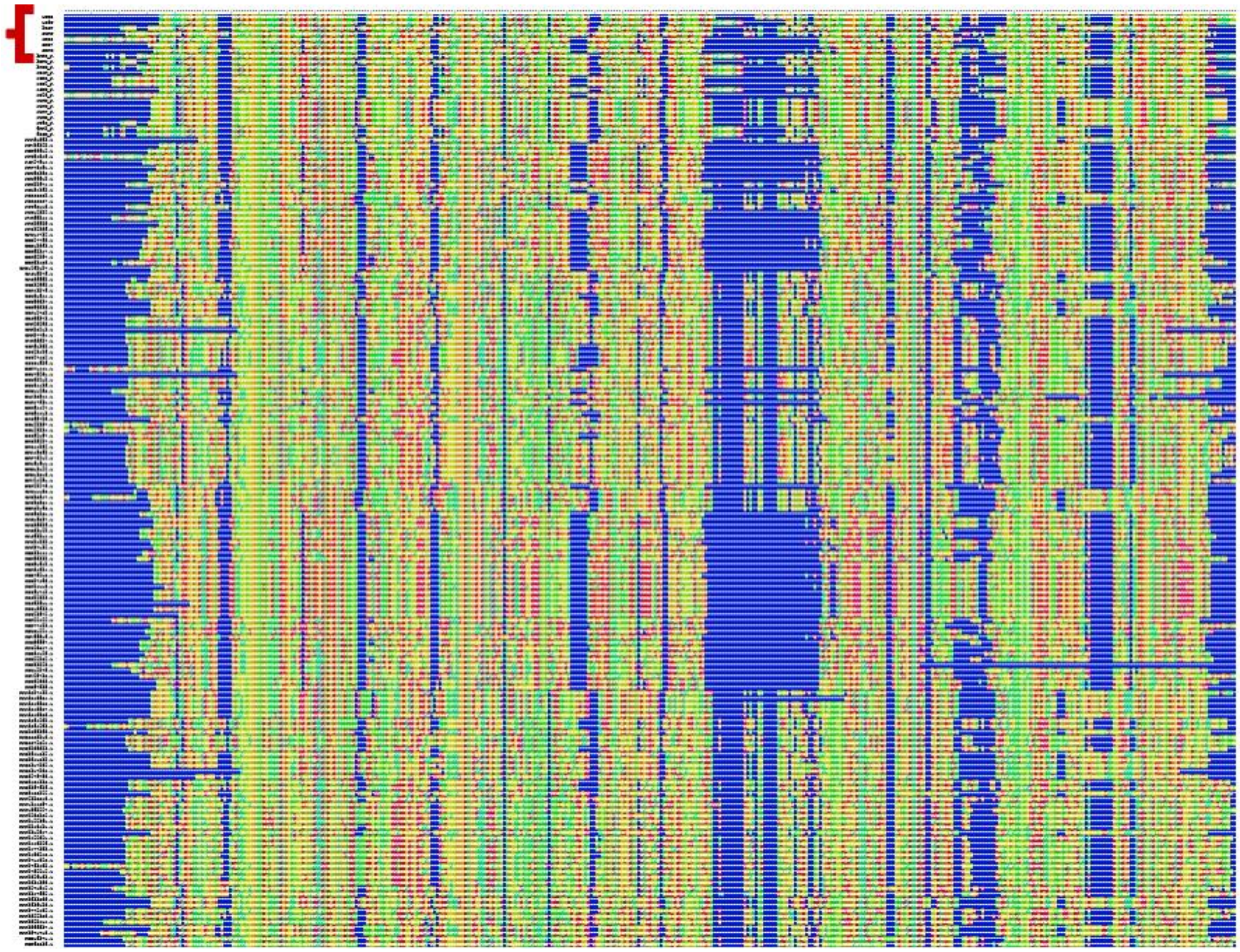
